# Information flow across the cortical timescales hierarchy during narrative construction

**DOI:** 10.1101/2021.12.01.470825

**Authors:** Claire H. C. Chang, Samuel A. Nastase, Uri Hasson

**Affiliations:** Princeton Neuroscience Institute, Princeton University, Princeton, New Jersey, 08540, USA

**Keywords:** cortical hierarchy, naturalistic stimuli, fMRI, functional connectivity, language processing

## Abstract

When listening to spoken narratives, we must integrate information over multiple, concurrent timescales, building up from words to sentences to paragraphs to a coherent narrative. Recent evidence suggests that the brain relies on a chain of hierarchically organized areas with increasing temporal receptive windows to process naturalistic narratives. We hypothesized that the structure of this cortical processing hierarchy should result in an observable sequence of response lags between networks comprising the hierarchy during narrative comprehension. This study uses functional MRI to estimate the response lags between functional networks during narrative comprehension. We use inter-subject cross-correlation analysis to capture network connectivity driven by the shared stimulus. We found a fixed temporal sequence of response lags—on the scale of several seconds—starting in early auditory areas, followed by language areas, the attention network, and lastly the default mode network. This gradient is consistent across eight distinct stories but absent in data acquired during rest or using a scrambled story stimulus, supporting our hypothesis that narrative construction gives rise to inter-network lags. Finally, we build a simple computational model for the neural dynamics underlying the construction of nested narrative features. Our simulations illustrate how the gradual accumulation of information within the boundaries of nested linguistic events, accompanied by increased activity at each level of the processing hierarchy, can give rise to the observed lag gradient.

**Significance Statement:** Our findings reveal a consistent, stimulus-driven gradient of lags in connectivity along the cortical processing hierarchy—from early auditory cortex to the language network, then to the default mode network—during the comprehension of naturalistic, spoken narratives. We provide a simple computational model for the neural dynamics underlying the construction of nested narrative features, allowing us to systematically explore the conditions under which the lag gradient emerges and synthesize our results with previous findings based on simple well-controlled language stimuli. Our results illustrate the isomorphism between hierarchically structured neural dynamics and hierarchically structured, real-world narrative inputs.

## Introduction

Narratives are composed of nested elements that must be continuously integrated to construct a meaningful whole, building up from words to phrases to sentences to a coherent narrative (1). Recent evidence suggests that the human brain relies on a chain of hierarchically organized brain areas with increasing temporal receptive windows (TRWs) to process this temporally evolving, nested structure (Fig. 1A). This cortical hierarchy was first revealed by studies manipulating the temporal coherence of naturalistic narratives (2, 3). These studies reported a topography of processing timescales where early auditory areas respond reliably to rapidly-evolving acoustic features, adjacent areas along the superior temporal gyrus respond reliably to information at the word level, and nearby language areas respond reliably only to coherent sentences. Finally, areas at the top of the processing hierarchy in the default mode network (DMN) integrate slower-evolving semantic information over many minutes (4).

**Fig. 1.**
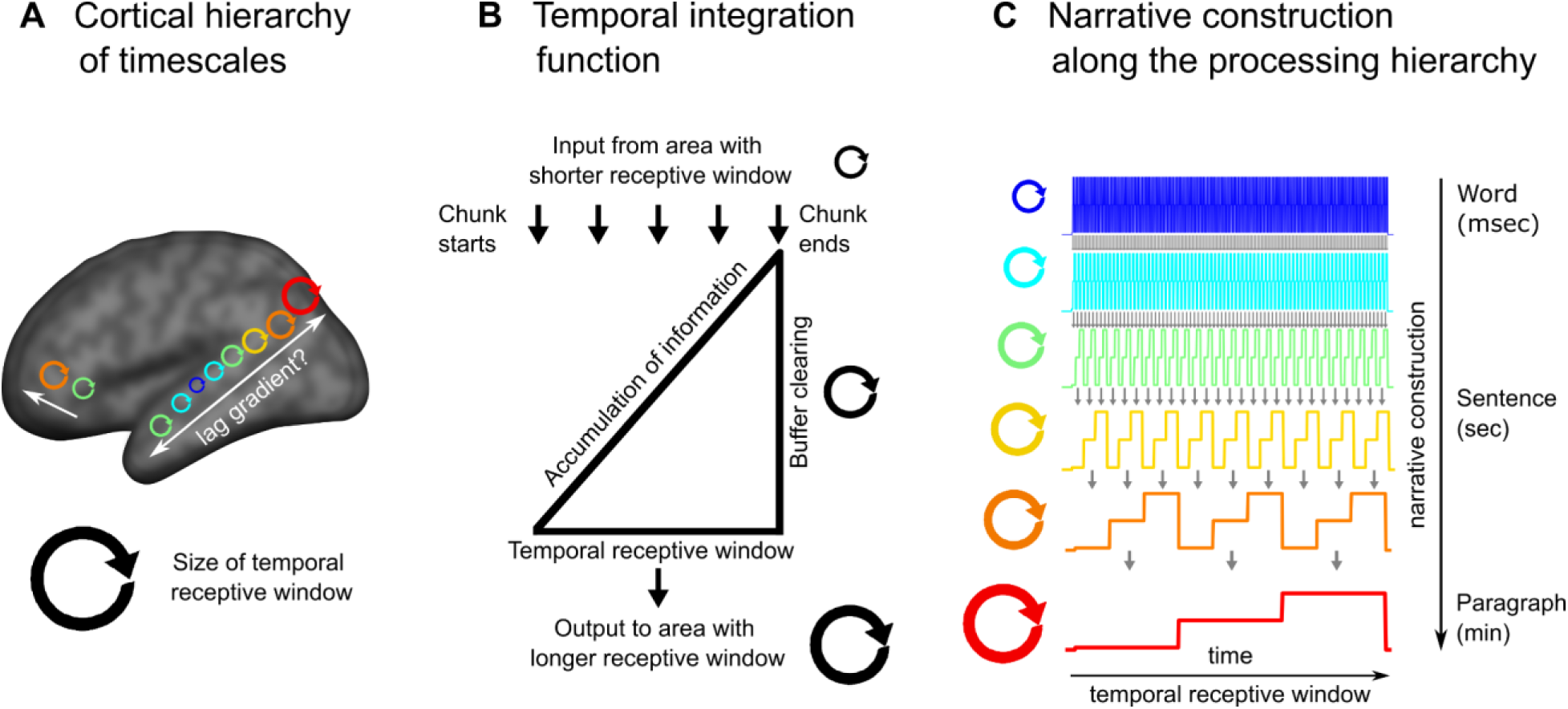
Narrative construction in the hierarchical processing framework. (A) The proposed cortical hierarchy of increasing temporal receptive windows (adapted from (5)). (B) Each level of the processing hierarchy continuously accumulates information over inputs from the preceding level. For example, phrases built over words are constructed into sentences. The accumulated information is flushed out at structural boundaries. (C) Each level of the processing hierarchy provides the building blocks for the next level, which naturally leads to longer temporal receptive windows, corresponding to linguistic units of increasing sizes. This model of narrative construction along the cortical processing hierarchy implies a gradient of response lags across the cortical hierarchy.

This cortical hierarchy of increasing temporal integration windows is thought to be a fundamental organizing principle of the brain (5, 6). The cortical hierarchy of TRWs in humans has been described using fMRI (2, 3, 7, 8) and ECoG (9). Recent work has shown that deep language models also learn a gradient or hierarchy of increasing TRWs (10–12), and that manipulating the temporal coherence of narrative input to a deep language model yields representations closely matching the cortical hierarchy of TRWs in the human brain (13). Furthermore, the cortical hierarchy of TRWs matches the intrinsic processing timescales observed during rest in humans (9, 14, 15) and monkeys (16). This cortical topography also coincides with anatomical and functional gradients such as long-range connectivity and local circuitry (17–19), which have been shown to yield varying TRWs (20, 21).

The proposal that the cortex is organized according to a hierarchy of increasing TRWs implies that each area “chunks” and integrates information at its preferred temporal window and that narrative construction proceeds along the cortical hierarchy. For example, an area that processes phrases receives information from areas that process words (Fig. 1B), which are further transmitted to areas that integrate phrases into sentences. At the end of each phrase, information is rapidly cleared to allow for real-time processing of the next phrase (1, 7). The chunking of information at varying granularity is supported by recent studies that used data-driven methods to detect boundaries as shifts between stable patterns of brain activity (22, 23).

This model of narrative construction (Fig. 1C) predicts a gradient of response lags across the cortical processing hierarchy; namely, shorter temporal lags among adjacent areas along the processing hierarchy than regions further apart in the cortical hierarchy. In the current study, we test this prediction by comparing response fluctuations elicited by spoken narratives in different brain areas using lag-correlation. We extract the lag with the peak correlation to estimate inter-region temporal difference. To focus on neural responses to linguistic and narrative information, we used inter-subject functional connectivity (ISFC) analysis (24, 25). Unlike traditional within-subject functional connectivity (WSFC), ISFC effectively filters out the idiosyncratic fluctuations that drive intrinsic functional correlations within subjects. Isolating stimulus-locked neural activity from intrinsic neural activity allows us for the first time to observe the temporal dynamics of narrative construction across the cortical hierarchy. We predicted that ISFC analysis would reveal an inter-region lag gradient during the comprehension of intact narrative, but not during scrambled-story or rest, which do not involve narrative construction. Finally, we provide a computational model to clearly illustrate how the construction of nested narrative features could give rise to the observed lag gradient, and how the lag gradient deteriorates without naturalistic inputs.

## Results

To test the hypothesis that narrative construction will yield a gradient of response lags across brain regions, we first divided the neural signals into six networks by applying k-means clustering to WSFC measured during rest (SI Appendix, Fig. S1). We labeled these networks based on anatomical correspondence with previously defined functional regions following Simony and colleagues (25), including the auditory (AUD), ventral language (vLAN), dorsal language (dLAN), default mode network (DMN), and attention (ATT) networks, aligning with the previously documented TRW hierarchy (SI Appendix, Fig. S2).

We computed lag-ISFC (i.e. cross-correlation) at varying temporal lags between all pairs of networks (Fig. 2A and SI Appendix, Fig. S3). The lags with maximum ISFC (i.e. “peak lag”) for each seed-target pair were extracted as an index for the temporal gaps in the stimulus-driven processing between each pair of networks. The extracted peak lags were color-coded to construct the network × network peak lag matrix (Fig. 2B and 2C). In the following, we describe the observed lag gradient in detail and several control analyses. Finally, we simulated the nested narrative structure and the corresponding brain responses to explore how different integration functions at different timescales could give rise to the observed lag gradient.

**Fig. 2.**
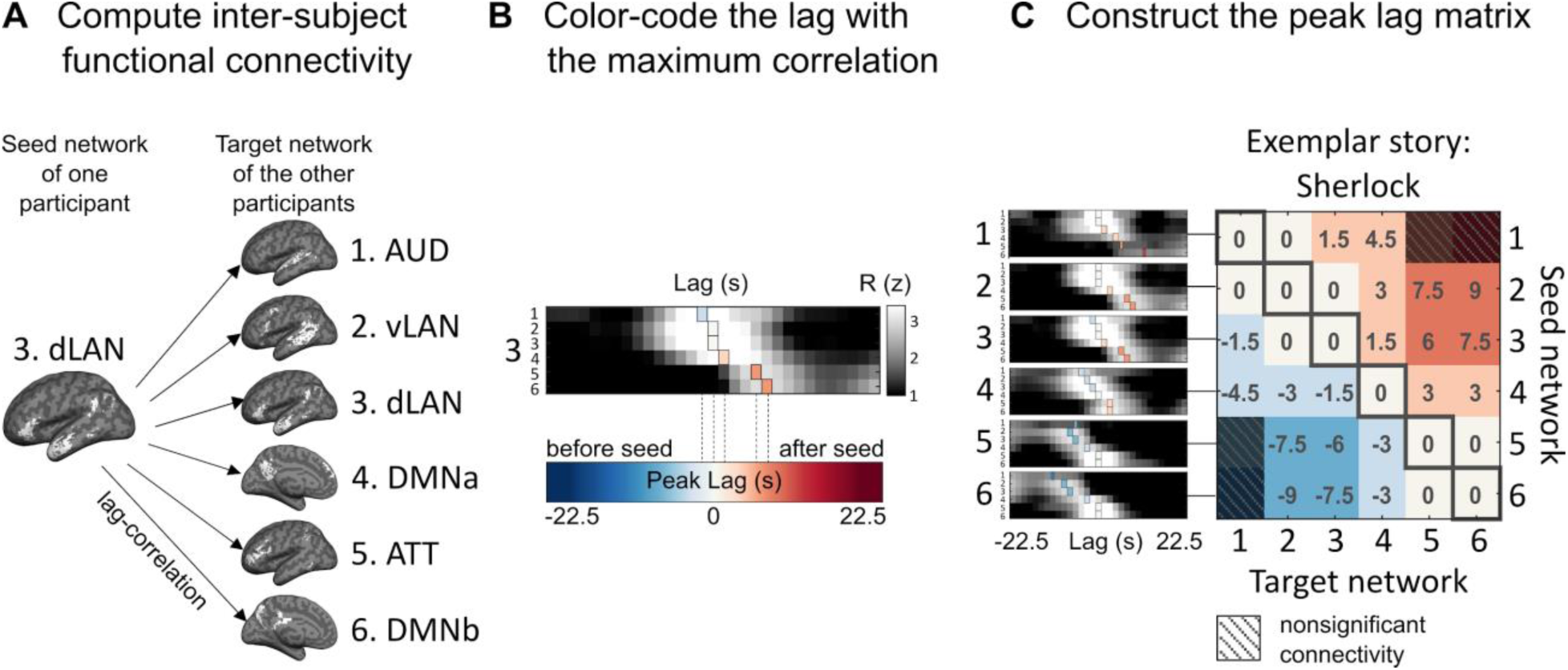
Construction of the inter-network peak lag matrix. (A) Lag-ISFC (cross-correlation) between seed-target network pairs were computed using the leave-one-subject out method. The dLAN network is used as an example seed network for illustrative purposes. (B) The matrix depicts ISFC between the dLAN seed and all six target networks at varying lags. The lag with the peak correlation value (colored vertical bars) was extracted and color-coded according to lag. For visualization, the lag-ISFCs were z-scored across lags. (C) The network × network peak lag matrix (p < .05, FDR corrected). Warm colors represent peak lags following the seed network, while cool colors represent peak lags preceding the seed network; zeros along the diagonal capture the intra-network ISC. An example story (“Sherlock”) is shown for illustrative purposes.

### Fixed lag gradient across cortical networks

The average lag-ISFC across stories was computed for each seed network (Fig. 3A, left). The lag-ISFC between a seed network and the same network in other subjects always peaked at lag 0, reflecting the strong stimulus-locked within-network synchronization reported in the ISC literature (3, 26, 27) (SI Appendix, Fig. S3). Interestingly, however, non-zero peak lags were found between different networks. Relative to a low-level seed, putatively higher-level networks showed peak connectivity at increasing lags. For example, the stimulus-induced activity in dLAN lagged 1 TR (1.5 s) behind activity in AUD, whereas the activity in DMNb lagged 4 TRs (6 s) behind activity in dLAN. Importantly, regardless of the choice of seed, the target networks showed peak connectivity in a fixed order progressing through AUD, vLAN, dLAN, DMNa, ATT, and DMNb.

**Fig. 3.**
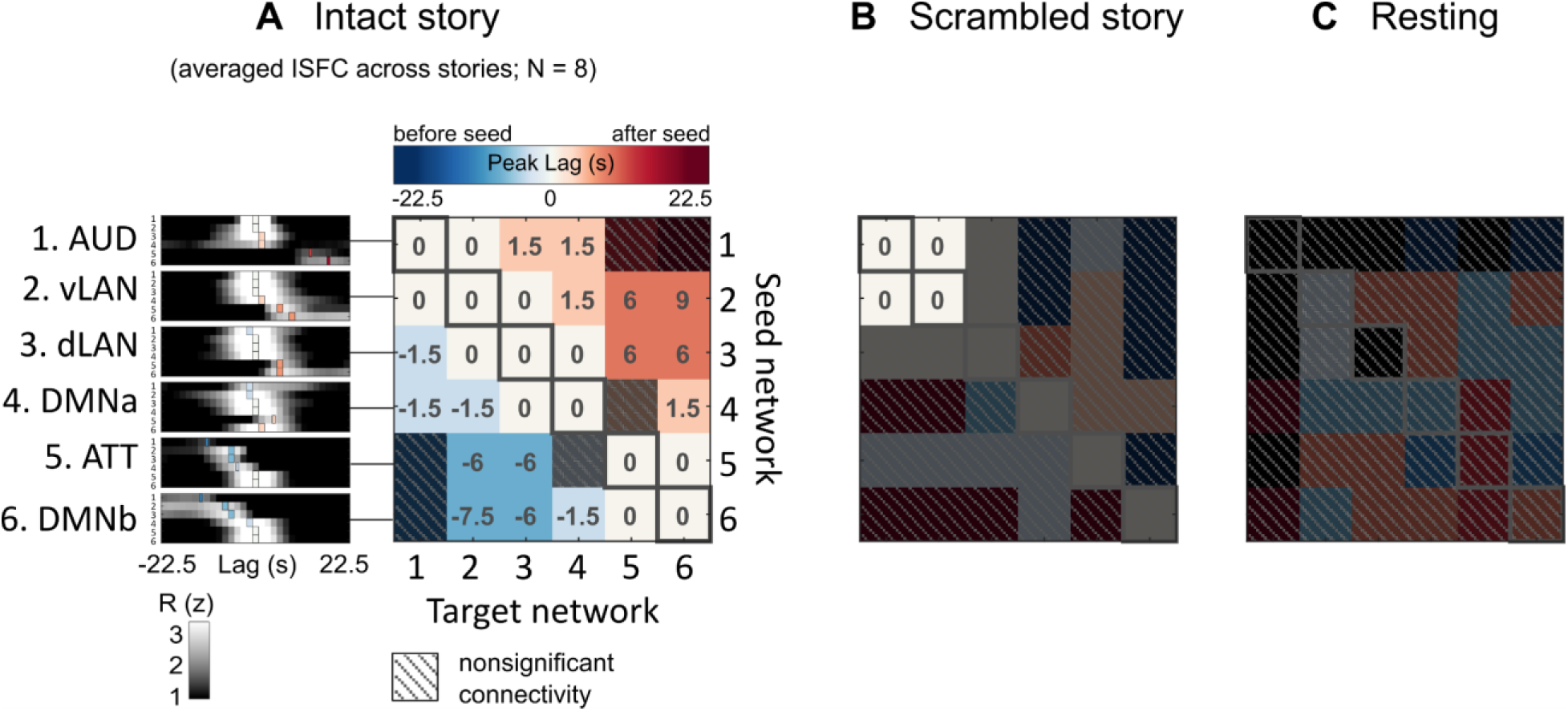
The peak lag matrix across eight stories reveals a fixed lag gradient across networks, which is abolished during scrambled narratives and rest. (A) The network × network peak lag matrix is based on the averaged lag-ISFC across eight stories. For visualization, lag-ISFC curves at left were z-scored across lags. (B) Peak lag matrix based on responses to a scrambled story stimulus (scrambled words). (C) Peak lag matrix based on resting-state data. Peak lag matrices are thresholded at p < .05 (FDR corrected).

To summarize the findings, we color-coded the peak lags and collated them into a peak lag matrix where each row corresponds to a seed network and each column corresponds to a target network (Fig. 3A, right; see SI Appendix, Fig. S4 for the lag-ISFC waveforms). The white diagonal indicates a peak at zero lag within each area, reflecting the intra-network synchronization across subjects (i.e. ISC) (SI Appendix, Fig. S3), while the cool-to-warm color gradient indicates a fixed order of peak lags. For example, the first row shows a white-to-warm gradient, reflecting that when AUD served as the seed, other networks were either synchronized with or followed AUD, but never preceded it. Conversely, the cool-to-white gradient of the last row indicates that all other networks preceded the DMNb seed. The lag gradient can also be observed in individual stories (SI Appendix, Fig. S5), although these patterns are noisier than the averaged results. The lag gradient proceeded in a fixed order across all networks, suggesting that bottom-up narrative construction is reflected in lagged connectivity between stages along the cortical hierarchy from AUD up to DMNb. Similar results were obtained when we defined the ROIs using the TRW hierarchy (SI Appendix, Fig. S2D).

### Temporal scrambling abolishes the lag gradient

We hypothesized that the lag gradient reflects the emergence of macroscopic story features (e.g. narrative situations or events) integrated over longer periods of time in higher-level cortical networks (22, 23). To support this point, we next used the same procedure to compute the peak lag matrix for a temporally scrambled version of one story (“Pie Man”; as for the results of intact “Pie Man”, please see *SI Appendix, Fig. S5*). In this dataset, the story stimulus was spliced at the word level and scrambled, thus maintaining similar low-level sensory statistics while abolishing the slower-evolving narrative content. The peak lag matrix for the scrambled story revealed synchronized responses at lag 0 both within and between the AUD and vLAN networks, but no significant peaks within or between other networks (Fig. 3B). This reflects low-level speech processing limited to the word level and indicates that disrupting the narrative structure of a story abolishes the temporal propagation of information to higher-level cortical areas.

### No lag gradient during rest

As an additional control, we also examined whether the lag gradient observed during the intact story could be detected during rest. The resting state is dominated by intrinsic fluctuations and there is no external stimulus to drive synchronized brain activity across subjects as well as propagation of activity across cortical areas. As expected, no significant ISFC peaks were found (Fig. 3B). This provides further evidence that the observed lag gradient is driven by the stimulus itself.

### Idiosyncratic within-subject fluctuations obscure the lag gradient

We next asked whether the inter-network lag gradient observed during spoken stories can be observed using traditional WSFC. As expected, WSFC analysis revealed a strong peak correlation at lag zero within each network, but also a peak correlation at lag zero across all networks such that no gradient was observed (SI Appendix, Fig. S6). This result supports the claim that ISFC analysis filters out intrinsic signal fluctuations that propagate across brain areas, revealing the propagation of shared story information across networks (24, 25). This result also verifies that intrinsic, inter-network differences in hemodynamic responses cannot account for the lag gradient; otherwise, WSFC should show a similar lag pattern as ISFC.

### Lag gradient across fine-grained subnetworks

To verify that the peak lag gradient could also be observed at a finer spatial scale, we further divided each of the six networks into ten subnetworks, again by applying k-means clustering to resting-state WSFC (k = 10 within each network). The peak lag matrix between the sixty subnetworks was generated using the same methods as in the network analysis (SI Appendix, Fig. S7A). We also visualized the brain map of lags between one selected seed (posterior superior/middle temporal gyrus) and all the target subnetworks (SI Appendix, Fig. S7B). Similar to the network level analysis, the peak lag between the subnetworks revealed a gradient from the early auditory cortex to the language network (auditory association cortex), then to the DMN.

### Dominant bottom-up lag gradient across networks

We adopted a method introduced by Mitra and colleagues (28) to discern whether there are multiple parallel lag sequences between networks. We applied principal component analysis (PCA) to the inter-network peak lag matrix (Fig. 3A) and examined the cumulative variance accounted for across principal components. Our results revealed that, at the coarse level of the cortical networks used here, the first principle component explains 88.8% of the variance in our lag matrix (SI Appendix, Fig. S8). This suggests that there is a single, unidirectional lag gradient across networks.

### The lag gradient is not driven by transient linguistic boundary effects

Prior work has reported that scene/situation boundaries in naturalistic stimuli elicit transient brain responses that vary across regions (29–34). To test whether this transient effect could drive the gradient observed in our lag matrix, we computed lag-ISFC after regressing out the effects of word, sentence, and paragraph boundaries in two stories with time-stamped annotations. As shown in SI Appendix, Fig. S9, the regression model successfully removed transient effects of the boundaries from the fMRI time series. Critically, however, the lag gradient remained qualitatively similar when accounting for boundaries, indicating that the observed lag gradient does not result from transient responses to linguistic boundaries in the story stimulus.

### Reproducing the lag gradient by simulating narrative construction

Narratives have a multi-level nested hierarchical structure (35) and are reported to elicit neural processing at increasingly long timescales along the cortical hierarchy (22, 23). To better understand how the construction of nested narrative features could give rise to the long inter-network lag gradient we observed, with up to 9-second lags, we created a simulation capturing the hierarchically nested temporal structure of real-world narratives and the corresponding hierarchy of cortical responses.

To match the six networks discussed so far, we simulated story features emerging across six distinct timescales, which roughly correspond to words, phrases, sentences, 2–3 sentences, and paragraphs. The initial level of the simulated narrative hierarchy was populated with relatively brief low-level units, with boundary intervals sampled from actual word durations in a spoken story (SI Appendix, Fig. S10). These simulated “words” were integrated into “phrases” of varying lengths with a mean length of three words to obtain second-level boundaries (Fig. 4A). All “phrase”-level boundaries were also “word”-level boundaries. A six-level structure was ultimately generated by recursively applying this procedure. Since paragraphs are often separated in real stories by longer silent periods (SI Appendix, Fig. S11), we inserted pauses at top-level (sixth-level) boundaries. The bottom-up construction of narrative structure gives rise to inter-level alignment and increasing processing timescales at higher levels, as proposed in the hierarchical processing framework (5, 6, 18).

**Fig. 4.**
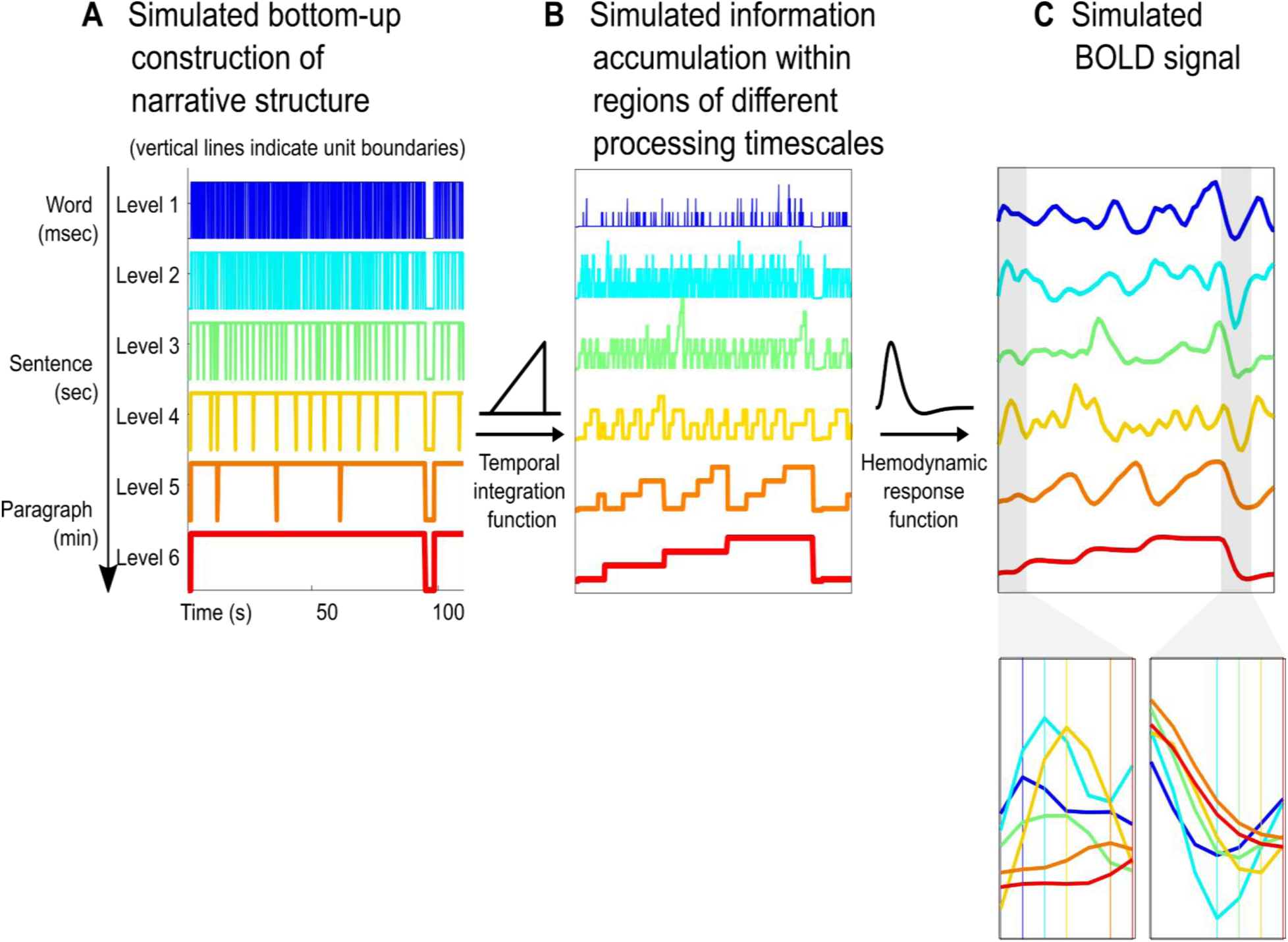
Simulating narrative construction and the corresponding brain responses. (A) The construction of the nested narrative structure, simulated by sampling boundary intervals from actual word durations and recursively integrating them to obtain structural boundaries at higher levels. (B) Information accumulation at different levels is generated by a linearly increasing temporal integration function. We postulated that information accumulation is accompanied by increased activity. (C) BOLD responses generated by HRF convolution. This visualization is based on parameters estimated from a spoken story stimulus (Table S1).

The simulated response amplitudes were generated using a linearly increasing temporal integration function (Fig. 4B), based on prior work showing that information accumulation is accompanied by gradually increasing activation within phrases/sentences (36–41) and paragraphs (29, 32) (a similar sentence/paragraph length effect was also observed in our data; see SI Appendix, Fig. S12). The linearly increasing temporal integration function accumulates activity derived from lower-level units within the interval between unit boundaries at the current levels and flushes out the accumulated activity at unit boundaries of the current level. To account for hemodynamic lag in the fMRI signal, we applied a canonical hemodynamic response function (HRF) to the simulated response amplitudes (Fig. 4C). We averaged the inter-level lag correlations across thirty different simulated structures (equivalent to 30 different stories) and extracted the peak lags. This peak-lag analysis parallels the analysis previously applied to the fMRI data.

The simulation allows us to systematically manipulate the narrative structure and the temporal integration function to reveal the conditions under which the lag gradient emerges. We first performed the simulation with a set of “natural” parameters roughly motivated by the temporal properties of our narrative stimuli and a simple temporal integration function reflecting linear temporal accumulation (Table S1).

This simple simulation is sufficient to reproduce the inter-network lag gradient observed in the fMRI data (Fig. 5A; as well as the ISFC at lag zero; SI Appendix, Fig. S13). In addition, we also compared the spectral properties of the simulated and real BOLD signals (SI Appendix, Fig. S14). We first computed the average power spectral density (PSD) across stories. Replicating results reported by Stephens and colleagues (15), we found stronger low-frequency fluctuations in regions with longer TRWs. Computing the PSD of the simulated brain responses similarly revealed increased low-frequency power in responses to high-level structures with longer intervals between boundaries. We then adjusted one parameter at a time to explore the parameter space constrained by natural speech.

**Fig. 5.**
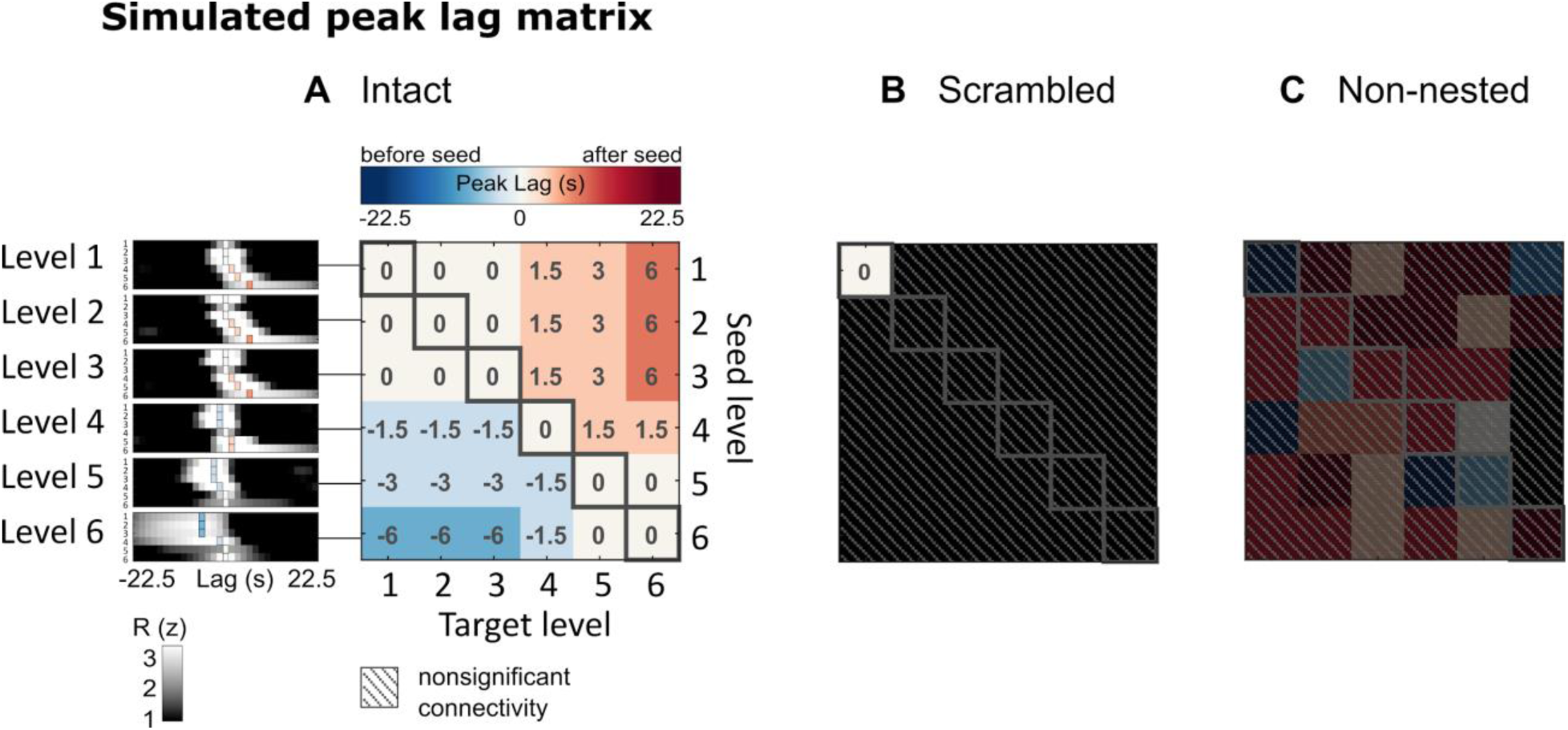
Simulated peak lag matrix. (A) Simulating the peak lag matrix observed during story-listening fMRI data (Fig. 3A) using parameters derived from a story stimulus (the same parameters as in Fig. 4 and Table S1). (B) Simulating the lag matrix observed during scrambled story (scrambled words) (Fig. 3B), by setting mean unit length = 1 and unit length variance = 0. (C) Lag matrix from the non-nested structure, created by combining levels extracted from independently generated nested structures, which disrupts the nesting relationship between different levels, similar to the scrambled story, while preserving the spectral properties of individual time series (p < .05, FDR correction).

### Key parameters for the emergence of a lag gradient

Within the bounds of natural speech (SI Appendix, Fig. S15), we observed that the simulated inter-network lag gradient is robust to varying lengths of linguistic/narrative units (mean: 2–4; variance: 0.1–1; longer length generated longer units, often with the top layers exceeding the length of the simulated story, i.e. 3000 words). The duration of inter-paragraph pauses was estimated from two stories (“Sherlock” and “Merlin”; SI Appendix, Fig. S11) (mean length: 1.5–4.5 sec; pause effect size: 0.01–1 SD of simulated activity). We also found that the model, similar to neural responses as observed by Lerner and colleagues (42), was robust to variations in speech rate (0.5– 1.5, relative to “Sherlock” speech rate). However, the lag gradient deteriorates with parameters outside of the bounds of natural speech, for example, when the inter-paragraph pause is set to 0 sec. We also simulated brain responses to word-scrambled stories by setting mean unit length = 1 and unit length variance = 0. With this setting, word-level units are never integrated into larger units (the units at each level correspond to individual words from the first level). No information integration is involved, resulting in flat activations and eliminating the difference in spectral properties of time series from different levels. No lag gradient is observed in this case (Fig. 5B).

Next, we computed inter-level lag-correlation using simulated responses to different nested structures (similar to responses to different stories), which preserves the spectral properties of individual time series while disrupting their nesting relationship. No significant lag-correlation was found when violating the nested structure of naturalistic narrative (Fig. 5C). In addition to the aforementioned linearly increasing integration function, we also explored several other temporal integration functions. We found that linearly and logarithmically increasing functions both yielded the inter-network lag gradient, but not the symmetric triangular or boxcar functions. The linearly decreasing function resulted in a reversed lag gradient (Fig. 6). These results suggest that the hierarchically nested structure that naturally arises from bottom-up narrative construction and a monotonically increasing integration function are key to the emergence of the lag gradient.

**Fig. 6.**
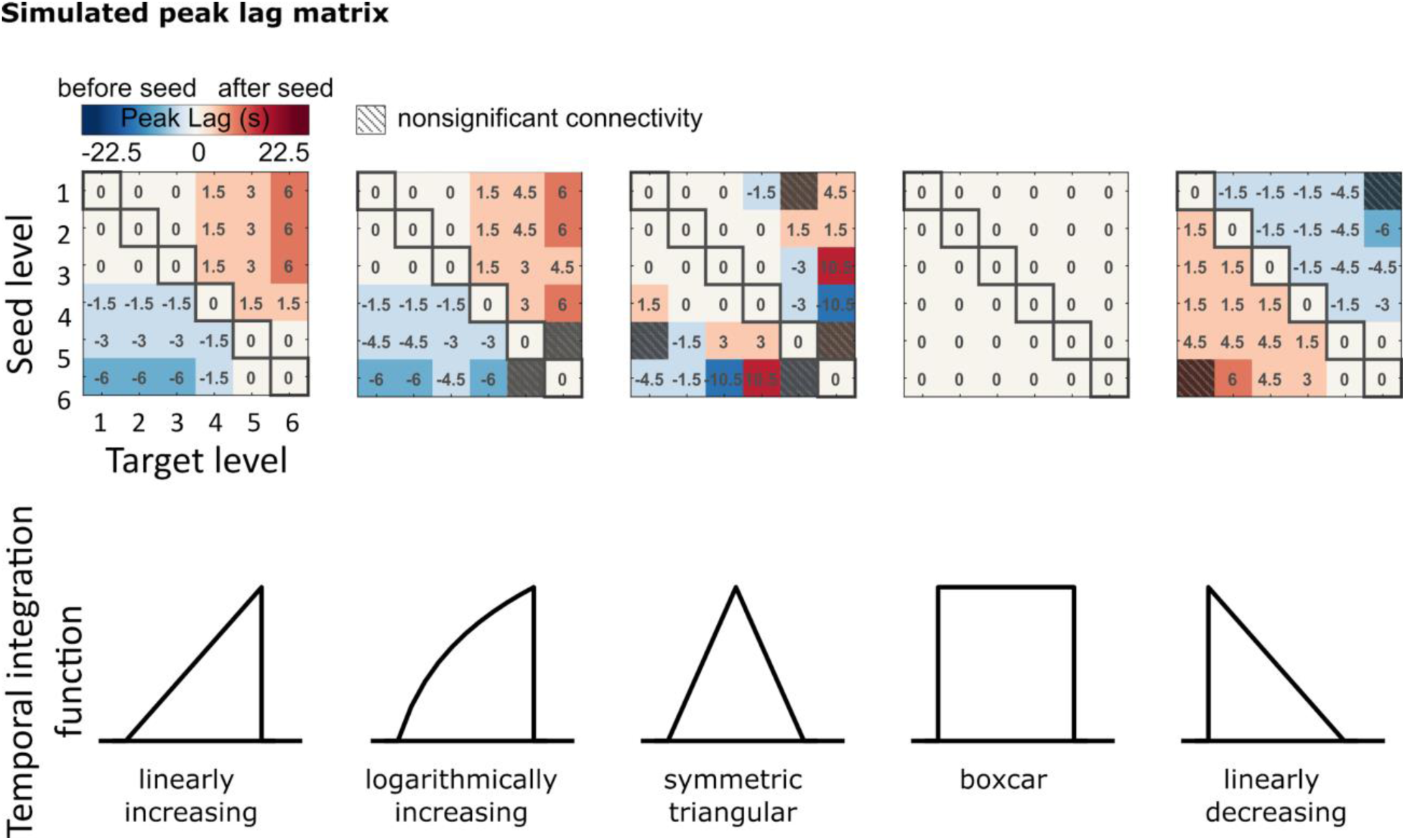
Lag matrices generated using different temporal integration functions (p < .05, FDR correction). The linearly and logarithmically increasing temporal integration functions yield a simulated peak lag matrix similar to the one observed in fMRI data; the symmetric triangle and boxcar functions, as well as the linearly decreasing function, do not.

## Discussion

By applying lag-ISFC to a collection of fMRI datasets acquired while subjects listened to spoken stories, we revealed a temporal progression of story-driven brain activity along a cortical hierarchy for narrative comprehension (Fig. 3A). The temporal cascade of cortical responses summarized by the inter-network lag gradient was consistent across stories as well as at coarse- and fine-grained functional network definitions (SI Appendix, Fig. S7). The results are in line with the hierarchical processing framework, which proposes a gradual emergence of narrative features of increasing duration and complexity along the processing hierarchy, from early sensory areas into higher-order cortical areas (Fig. 1). In support of our interpretation, we found that the lag gradient is absent during rest when there is no stimulus-evoked processing (Fig. 3B), and also when the temporal structure of the story is disrupted due to word scrambling (Fig. 3C).

We observed inter-network lags on the scale of several seconds (up to 9 seconds), reflecting the temporal structure of real-world narratives, which integrate sounds into words, sentences, and ideas over many seconds. Such long lags cannot be explained by regional variations in neurovascular coupling (43) or transient activity impulses at event boundaries. If the lag gradient only reflects variations in neurovascular coupling across regions, it should be present both when we isolate stimulus-driven activity using ISFC and when we examine idiosyncratic neural responses using WSFC. Instead, however, the lag gradient was detected only with ISFC, but not WSFC (SI Appendix, Fig. S6). Furthermore, differences in the hemodynamic response function across brain areas are usually reported at shorter timescales (e.g. ∼1–2 seconds) (44, 45) than the 0–9-second inter-network lag differences observed here in the context of narrative comprehension (Fig. 3). In addition, we found that transient event boundaries (29–34) did not account for the lag gradient (SI Appendix, Fig. S9).

Our simulation illustrates how narrative construction can give rise to the inter-network lag gradient by identifying three necessary conditions: (a) a cortical hierarchy of increasing processing timescales (Fig. 1 & Fig. 4) (5); (b) hierarchically nested linguistic/narrative events of increasing size along the processing hierarchy (Fig. 5B & 5C) (22, 23); and (c) gradual increasing brain activity, along with information accumulation, within the boundaries of events at each processing level (29, 32, 36–41), combined with a reset of activity (buffer clearing) at event boundaries (7) (see temporal integration function in Fig. 1B and Fig. 6). In this simple model, information integration at varying granularity (e.g. word, sentence, and paragraph) is sufficient to yield the inter-network lag gradient (Fig. 5) and spectral properties observed in the fMRI data (SI Appendix, Fig. S14). Minor adjustments to other parameters within the bounds of natural speech (i.e. speech rate, silent pause, and length of linguistic/narrative unit) did not change the gradient pattern (SI Appendix, Fig. S15).

The simulation provides a simple model which bridges the discovery of TRWs using natural stimuli (2, 42) and the accumulation of activity within linguistic units found using simple, well-controlled stimuli (e.g. sentences with similar structures) (36–41). Importantly, we note that our model is not the only one that could generate the predicted lag gradient. Our aim is to combine separate findings that point to the same cortical hierarchy with the simplest model possible. In addition, narrative processing is unlikely to be purely unidirectional (46). The lag gradient only captures the dominant process of bottom-up narrative construction (SI Appendix, Fig. S8). More studies are needed to examine recurrent or bidirectional connectivity, causal relations between networks, and nonstationary information flow over time.

Our results are also consistent with reports on the spatiotemporal dynamics of brain responses to naturalistic stimuli. A hierarchically nested spatial activation pattern has been revealed using movie, spoken story, and music stimuli (22, 23, 47). Chien and colleagues (7) reported a gradual alignment of context-specific spatial activation patterns, which was rapidly flushed at event boundaries, similar to the temporal integration function we adopted here. Taken together, the empirical findings, combined with our simulation, indicate that the spatiotemporal neural dynamics reflect the structure of naturalistic, ecologically-relevant inputs (6) and that such information is preserved even with the poor temporal resolution of fMRI. Although the current findings are derived from listener-listener coupling, the inter-regional dynamics may shed light on the lags observed in speaking-listener coupling (48–53). Given a particular seed region in the speaker’s brain, we would expect to observe coupling at differing lags for different target regions in the listener’s brain, and these lags may vary based on the temporal structure of the speaker’s narrative.

Our results demonstrate both the importance of using inter-subject methods to isolate stimulus-driven signals and the value of data aggregation. The fact that we obtained non-zero inter-network lag only with ISFC but not WSFC (SI Appendix, Fig. S6) indicates that stimulus-driven network configuration may be masked by the idiosyncratic fluctuations that dominate WSFC analyses (24, 25). Furthermore, although the inter-network lags could be observed within individual stories (SI Appendix, Fig. S5), the gradient pattern is much clearer after aggregating across stories (Fig. 3). Data aggregation is particularly important when using naturalistic stimuli because it is impossible to control the structure of each narrative (e.g. speaking style, duration, complexity, and content) (35, 54–56, 56). With these methods, we are able to reveal the inter-network lag gradient driven by naturalistic narratives, as predicted by the model shared information flow along the cortical processing hierarchy. Further work will be needed to examine recurrent or bidirectional information flow and to decode the content of narrative representations—specific to each story—as they are transformed along the cortical hierarchy.

## Materials and Methods

### fMRI datasets

This study relied on eight openly available spoken story datasets. Seven datasets were used from the “Narratives” collection (OpenNeuro: https://openneuro.org/datasets/ds002245) (57), including “Sherlock” and “Merlin” (18 participants, 11 females) (52), “The 21^st^ year” (25 participants, 14 females) (58), “Pie Man (PNI)”, “I Knew You Were Black”, “The Man Who Forgot Ray Bradbury”, and “Running from the Bronx (PNI)” (48 participants, 34 females). One dataset was used from Princeton Dataspace: “Pie Man” (36 participants, 25 females) (https://dataspace.princeton.edu/jspui/handle/88435/dsp015d86p269k) (25).

Two non-story datasets were also included as controls: a word-scrambled “Pie Man” (36, participants, 20 females) dataset and a resting-state dataset (36 participants, 15 females) (see the Princeton DataSpace URL above) (25).

All participants reported fluency in English and were 18–40 years in age. The criteria of participant exclusion have been described in previous studies for “Sherlock”, “Merlin”, “The 21^st^ year”, and “Pie Man.” For “Pie Man (PNI)”, “I Knew You Were Black”, “The Man Who Forgot Ray Bradbury”, and “Running from the Bronx (PNI),” participants with comprehension scores 1.5 standard deviations lower than the group means were excluded. One participant was excluded from “Pie Man (PNI)” for excessive movement (translation along the z-axis exceeding 3 mm).

All participants provided informed, written consent, and the experimental protocol was approved by the institutional review board of Princeton University.

### fMRI preprocessing

fMRI data were preprocessed using FSL (https://fsl.fmrib.ox.ac.uk/), including slice time correction, motion correction, and high-pass filtering (140 s cutoff). All data were aligned to standard 3 × 3 × 4 mm Montreal Neurological Institute space (MNI152). A gray matter mask was applied.

### Functional networks

Following Simony and colleagues (25), we defined 6 intrinsic connectivity networks within regions showing reliable responses to spoken stories. Voxels showing top 30% ISC in at least 6 out of the 8 stories were included. Using the k-means method (L1 distance measure), these voxels were clustered according to their group-averaged within-subject functional connectivity with all the voxels during resting. We refer to these functional networks as the auditory (AUD), ventral language (vLAN), dorsal language (dLAN), attention (ATT), and default mode (DMNa and DMNb) networks (SI Appendix, Fig. S1A). To ensure that our results hold for finer-grained functional networks, we further divided each of the six networks into ten subnetworks, again by applying k-means clustering to resting-state WSFC (k=10 within each superordinate network).

To compare these intrinsic functional networks to the TRW hierarchy, we computed the TRW index (i.e. intact > word-scrambled story ISC) following (8) for voxels within regions showing reliable responses to spoken stories, using the intact and word-scrambled Pie Man. Six TRW networks were then generated by splitting the TRW indices into six bins by five quantiles (SI Appendix, Fig. S2).

### WSFC, ISFC, and ISC

In this study, within-subject functional connectivity (WSFC) refers to within-subject inter-region correlation, while inter-subject functional connectivity (ISFC) refers to inter-subject inter-region correlation. Inter-subject correlation (ISC) refers to a subset of ISFC, namely, ISFC between homologous regions (SI Appendix, Fig. S3). ISFC and ISC were computed using the leave-one-subject-out method, i.e. correlation between the time series from each subject and the average time series of all the other subjects (24).

Before computing the correlation, the first 25 and last 20 volumes of fMRI data were discarded to remove large signal fluctuations at the beginning and end of time course due to signal stabilization and stimulus onset/offset. We then averaged voxelwise time series across voxels within network/region masks and z-scored the resulting time series.

Lag-correlations were computed by circularly shifting the time series such that the non-overlapping edge of the shifted time series was concatenated to the beginning or end. The left-out subject was shifted while the average time series of the other subjects remained stationary. Fisher’s z transformation was applied to the resulting correlation values prior to further statistical analysis.

### ISFC lag matrix

We computed the network × network × lag-ISFC matrix (SI Appendix, Fig. S3) and extracted the lag with peak ISFC (correlation) value for each network pair (Fig. 2). The peak ISFC value was defined as the maximal ISFC value within the window of lags from -15 to +15 TRs; we required that the peak ISFC be larger than the absolute value of any negative peak and excluded any peaks occurring at the edge of the window.

To obtain the mean ISFC across stories, we applied two statistical tests. Only ISFC that passed both tests were considered significant. First, we performed a parametric one-tailed one-sample t-test to compare the mean ISFC against zero (N = 8 stories) and corrected for multiple comparisons by controlling the false discovery rate (FDR; (59); 6 seed × 6 target × 31 lags; q < .05).

Second, to exclude ISFC peaks that only reflected shared spectral properties, we generated surrogates with the same mean and autocorrelation as the original time series by time-shifting and time-reversing. We computed the correlation between the original seed and time-reversed target with time-shifts of -100 to +100 TRs. The resulting ISFC values were averaged across stories and served as a null distribution. A one-tailed z-test was applied to compare ISFCs within the window of lag -15 to +15 TRs against this null distribution. The FDR method was used to control for multiple comparisons (seed × target × lags; q < .05). When assessing ISFC for each story, only this second test was applied and all possible time-shifts were used to generate the null distribution.

### Principal component analysis of the lag matrix

We examined whether multiple lag sequences similarly contributed to the lag matrix, using the method introduced by Mitra and colleagues (28). We applied PCA to the lag matrix obtained from the averaged ISFC across stories, after transposing the matrix and zero-centering each column. Each principal component represents a pattern of relative lags, in other words, lag sequences. We computed the proportion of overall variance in the lag matrix accounted for by each component in order to determine whether more than one component played an important role.

### Word/sentence/paragraph boundary effect

To test the transient effect of linguistic boundaries on inter-network lag, we computed the lag-ISFC after regressing out activity impulses at boundaries. A multiple regression model was built for each subject. The dependent variable was the averaged time series of each network, removing the first 25 scans and the last 20 scans as in the ISFC analysis. The regressors included an intercept, the audio envelope, and three sets of finite impulse functions (−5 to +15 TRs relative to boundary onset), corresponding to word, sentence, and paragraph (event) boundaries. We then recomputed lag-ISFC based on the residuals of the regression model.

### Word/sentence/paragraph length effect

We replicated the sentence length (36–41) and paragraph length (29, 32) effect with the “Sherlock” and “Merlin” datasets, which were collected from the same group of participants. The onsets and offsets of each word, sentence, and paragraph (event) were manually time-stamped. Given the low temporal resolution of fMRI (TR = 1.5 sec) and the difficulty of labeling the onset/offset of each syllable, they were estimated by dividing the duration of each word by the number of syllables it contains.

We built individual GLM models that included regressors corresponding to the presence of syllable, word, sentence, and paragraph respectively, accompanied by three parametric modulators: accumulated syllable number within words, accumulated word number within sentences, and accumulated sentence number within paragraphs. These parametric regressors were included to test whether brain activations accumulate toward the end of word/sentence/paragraph; the longer the word/sentence/paragraph the stronger the activations. In addition to the regressors of interest, one regressor was included for speech segments without clear paragraph labels. We did not orthogonalize the regressors to each other.

Effect maps of the three parametric modulators (i.e. word length, sentence length, and paragraph length) from the individual level models of both stories were smoothed with a Gaussian kernel (FWHM = 8 mm) and input to three group-level models to test the word, sentence, and paragraph length effects respectively (flexible factorial design including the main effects of story and participant; p < .005, not corrected). We observed sentence and paragraph length effects. Using the same threshold, no word length effect was observed,

### Power-spectral density analysis

We performed spectral analyses following (15). We estimated the power spectrum of the primary auditory area and a DMN area (precuneus). As for the connectivity analysis, we cropped the first 25 and last 20 scans and z-scored the time series. For each story, the resulting time series were averaged across subjects and normalized across time. The power spectrum of the group-mean time series was estimated using Welch’s method with a Hamming window of width 99 sec (66 TRs) and 50% overlap (based on the parameters from (15)). The power spectra of individual voxels were averaged within the anatomical masks of left Heschl’s gyrus and left precuneus from the AAL atlas. The mean spectra across stories were then computed.

### Simulating the construction of nested narrative structures and the corresponding BOLD responses

To illustrate how information accumulation at different timescales could account for the inter-network lag gradient during story-listening, we simulated the construction of nested narrative structures closely following the statistical structure of real spoken stories and generated BOLD responses at each processing level. To build the first level of a nested structure, we sampled a sequence of 3000 word durations with replacement from “Sherlock,” which is the longest example of spontaneous speech among our datasets, recorded from a non-professional speaker without rehearsal or script (SI Appendix, Fig. S10). Boundaries between units at the first level were set up accordingly.

### Unit length

First-level units were integrated into units of the next level with a lognormal distributed unit length; e.g. integrating three words into a phrase (unit length = 3) (SI Appendix, Fig. S10). Boundaries between second-level units were inserted accordingly. Second-level units were integrated into the third-level units following the same method. A nested structure of six levels was thus generated.

### Temporal integration function

Postulating that information accumulation is accompanied by increased activity, brain responses within each level of the nested structure were generated as a function of unit length. For example, a linear temporal integration function generates activity [1 2 3] for a “phrase” (i.e. a Level 2 unit) consisting of three “words” (i.e. Level 1 units). The first (word) level integration was computed based on syllable numbers sampled from “Sherlock” along with word durations.

### Pause length and pause effect size

In naturalistic narratives, boundaries between high-level units are often accompanied by silent pauses (SI Appendix, Fig. S11). Therefore, we inserted pauses with normally distributed lengths at the boundaries of the highest level units (SI Appendix, Fig. S10). Activity during the pause period was set as 0.1 standard deviations below the minimum activity of each level.

To account for HRF delay in fMRI signals, we applied the canonical hemodynamic response function provided by the software SPM (https://www.fil.ion.ucl.ac.uk/spm/) (60) and resampled the output time series from a temporal resolution of 0.001 sec to 1.5 sec to match the TR in our data. We ran 30 simulations for each set of simulation parameters. Each simulation produced different narrative structures (equivalent to different stories). The peak lag of the mean inter-level correlation across simulations was extracted and thresholded using the same method as in the ISFC analysis (Fig. 2).

To examine whether the simulated and real fMRI signals shared similar power spectra, we also applied the power-spectral density analysis to the simulated BOLD responses at each of the six levels and averaged across thirty simulations.

We started with a set of reasonable parameters (SI Appendix, Table 1) (speech rate = 1, relative to “Sherlock”; unit length mean = 3; unit length variance = 0.5; temporal integration function = linearly increasing; mean pause length = 3 sec; pause effect size = 0.1 SD of the simulated activity) and explored alternative parameter sets within the bound of natural speech to test whether inter-level lag was robust to parameter changes.

## Data availability

This study relied on eight openly available spoken story datasets. Seven datasets were used from the “Narratives” collection (OpenNeuro: https://openneuro.org/datasets/ds002245) (57), One dataset was used from Princeton Dataspace: “Pie Man” (36 participants, 25 females) (https://dataspace.princeton.edu/jspui/handle/88435/dsp015d86p269k) (25).

## Acknowledgment

This study is supported by the National Institute of Mental Health (R01-MH112357 and DP1-HD091948).

---

**Fig. S1.**
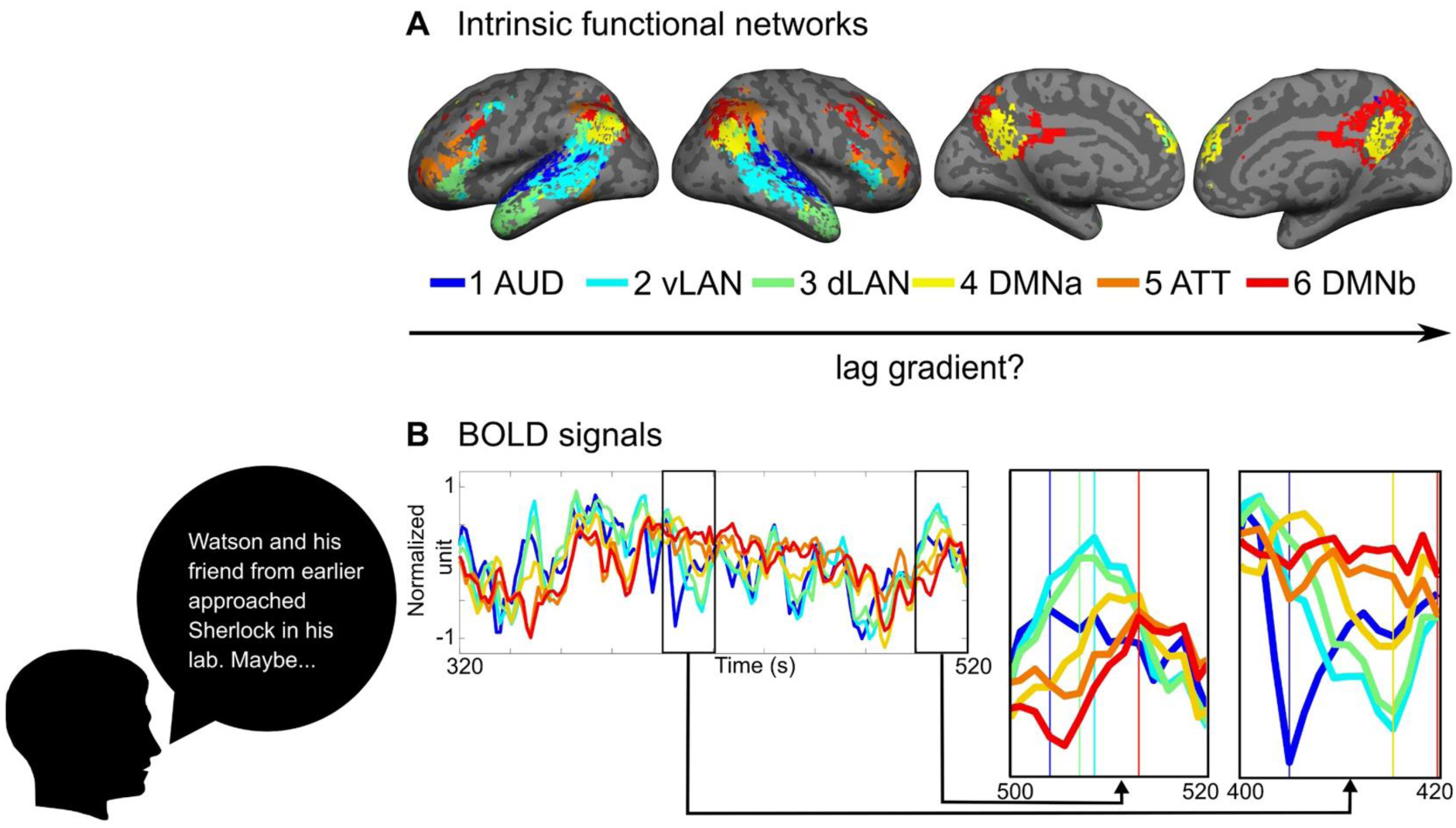
Averaged fMRI response time series for six intrinsic functional networks while subjects listened to a spoken story. (A) Functional networks defined by applying k-means clustering to WSFC measured during rest and labeled based on anatomical correspondence with previously defined functional regions following (25) (AUD: auditory; vLAN: ventral language; dLAN: dorsal language; DMN: default mode network; ATT: attention network). (B) Averaged fMRI responses time series in the “Sherlock” dataset, extracted from the predefined network masks. Two example segments of the response time series are highlighted at the bottom right. The peaks of the fluctuations in a given window are indicated by colored vertical lines. Note the stereotyped lag in both positive and negative BOLD fluctuations across networks; e.g. signal deflections in the dark blue auditory network tend to precede deflections in the cyan/green language networks, which tend to precede deflections in the yellow/red default mode networks.

**Fig. S2.**
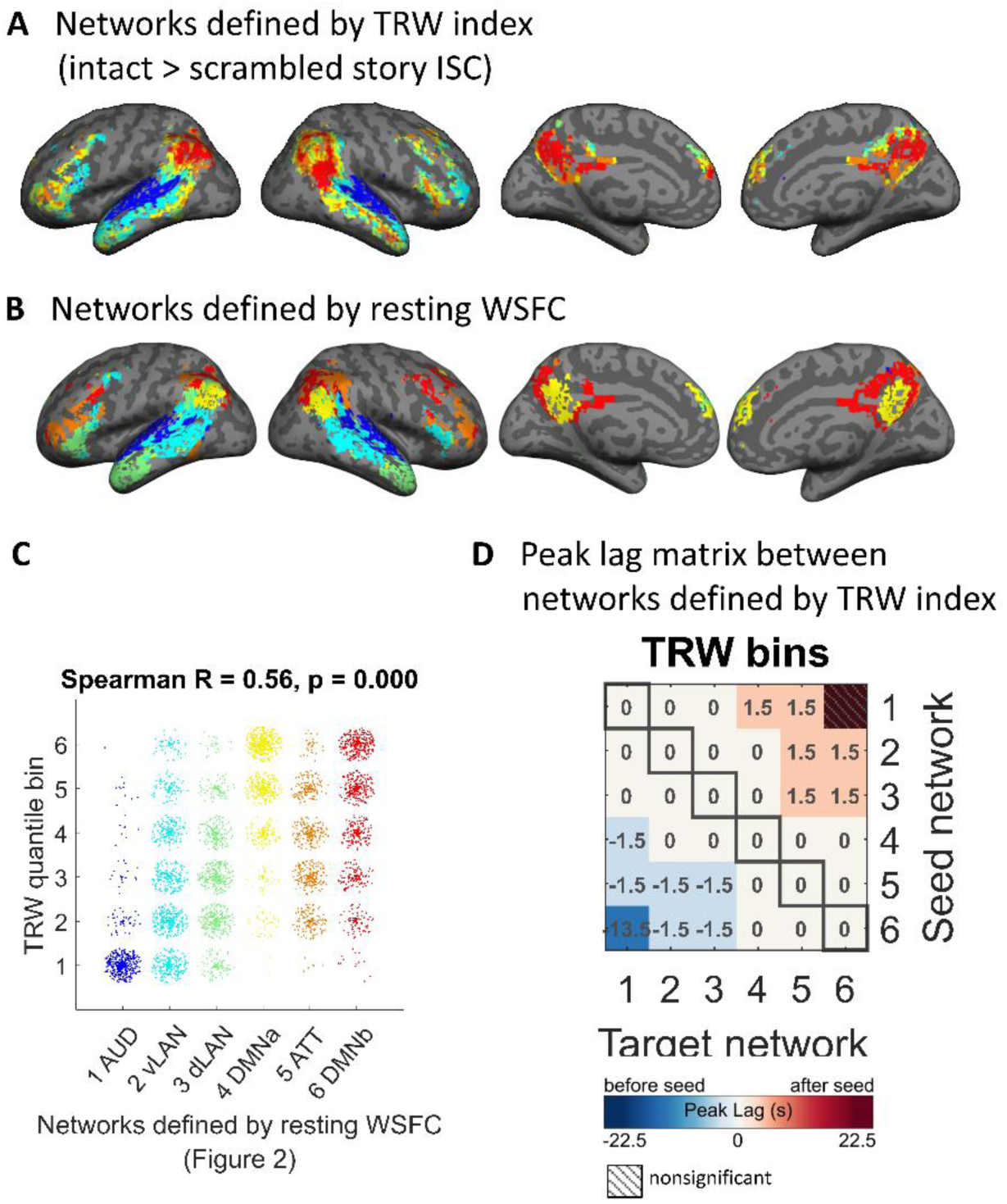
Lag gradient between networks defined by TRW indices. (A) Networks generated by splitting the TRW indices (intact > scrambled story ISC) into 6 bins by five quantiles. (B) networks defined by applying k-mean clustering to resting WSFC (SI Appendix, Fig. S1). (C) Networks defined by TRW index show a similar topographic gradient as the networks defined by resting-state WSFC, from the auditory areas to DMN, which is manifested by the significant correlation between the two sets of networks index. Random jitters are added to better show the overlapped data points. (D) Peak lag matrix between networks defined by TRW index across seven stories (p < .05, FDR corrected). “Pie Man” was excluded from this analysis since it was used to compute the TRW index.

**Fig. S3.**
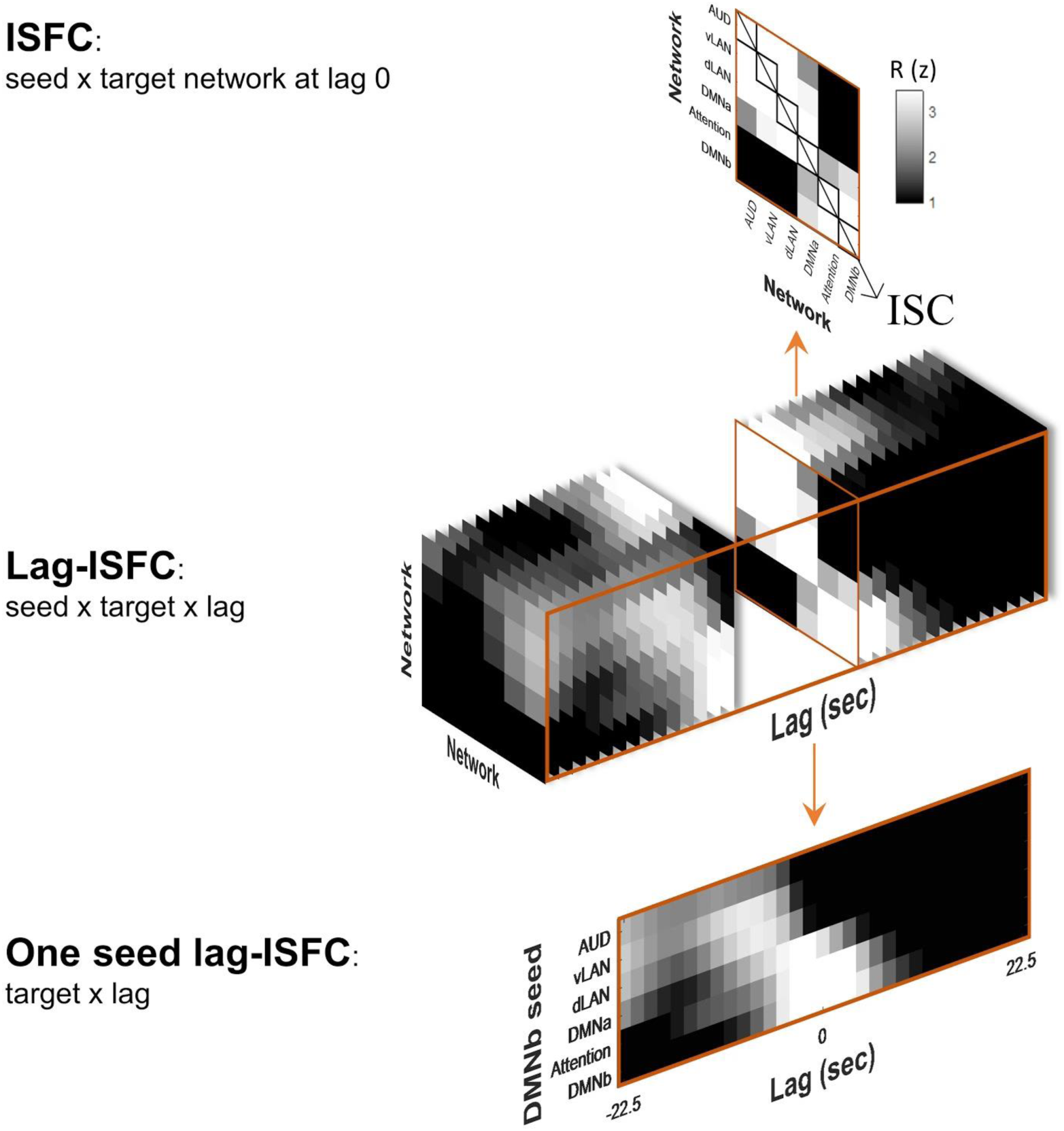
The relationship between inter-subject functional connectivity (ISFC), inter-subject-correlation (ISC), and lag-ISFC. This figure shows real data from the “Sherlock’’ story.

**Fig. S4.**
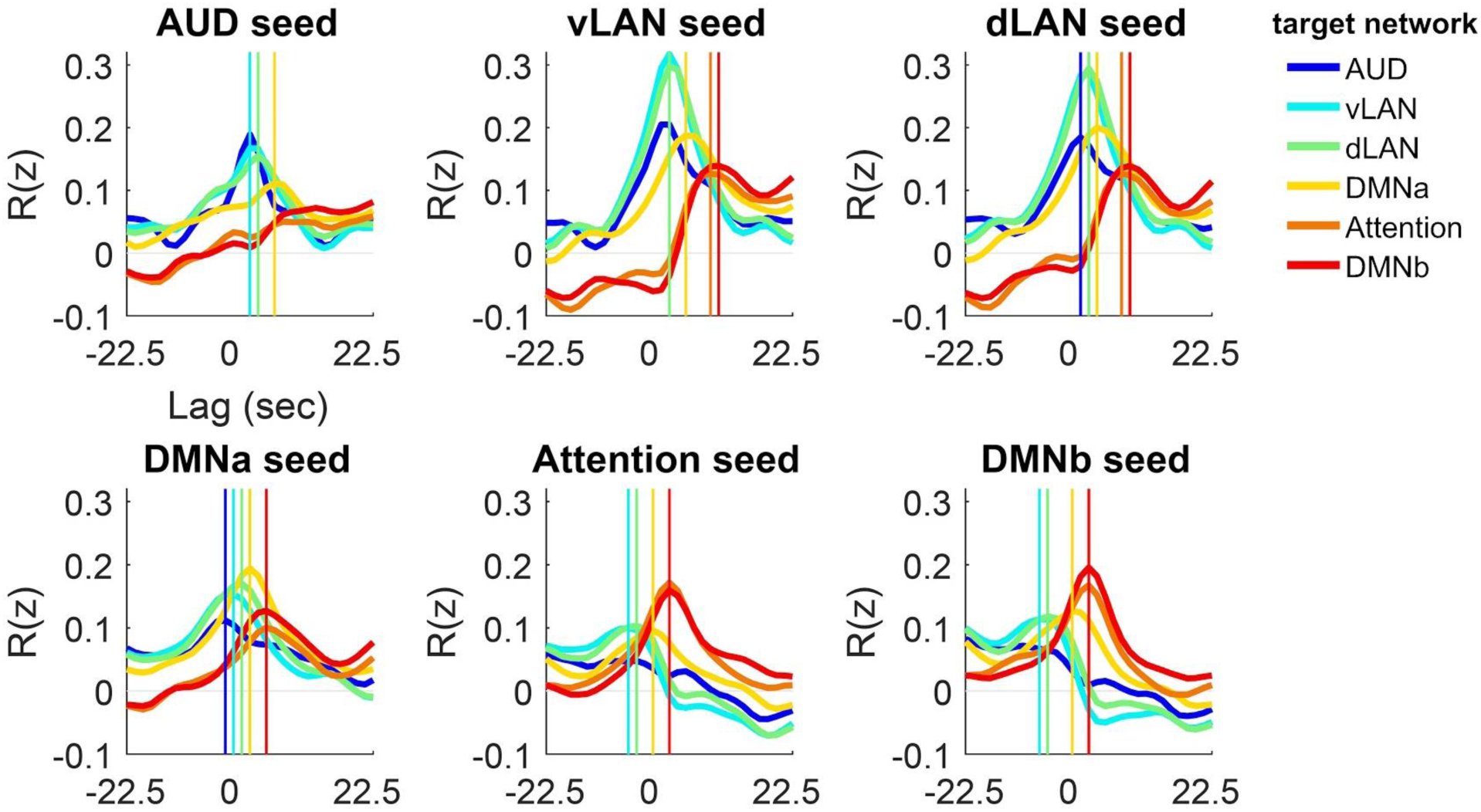
Lag-ISFC curves between the six functional networks, corresponding to the left panel of Fig. 3A. Fisher’s z transformation was applied to the R-values before averaging. Vertical lines indicate significant R peaks (p < .05, FDR correction).

**Fig. S5.**
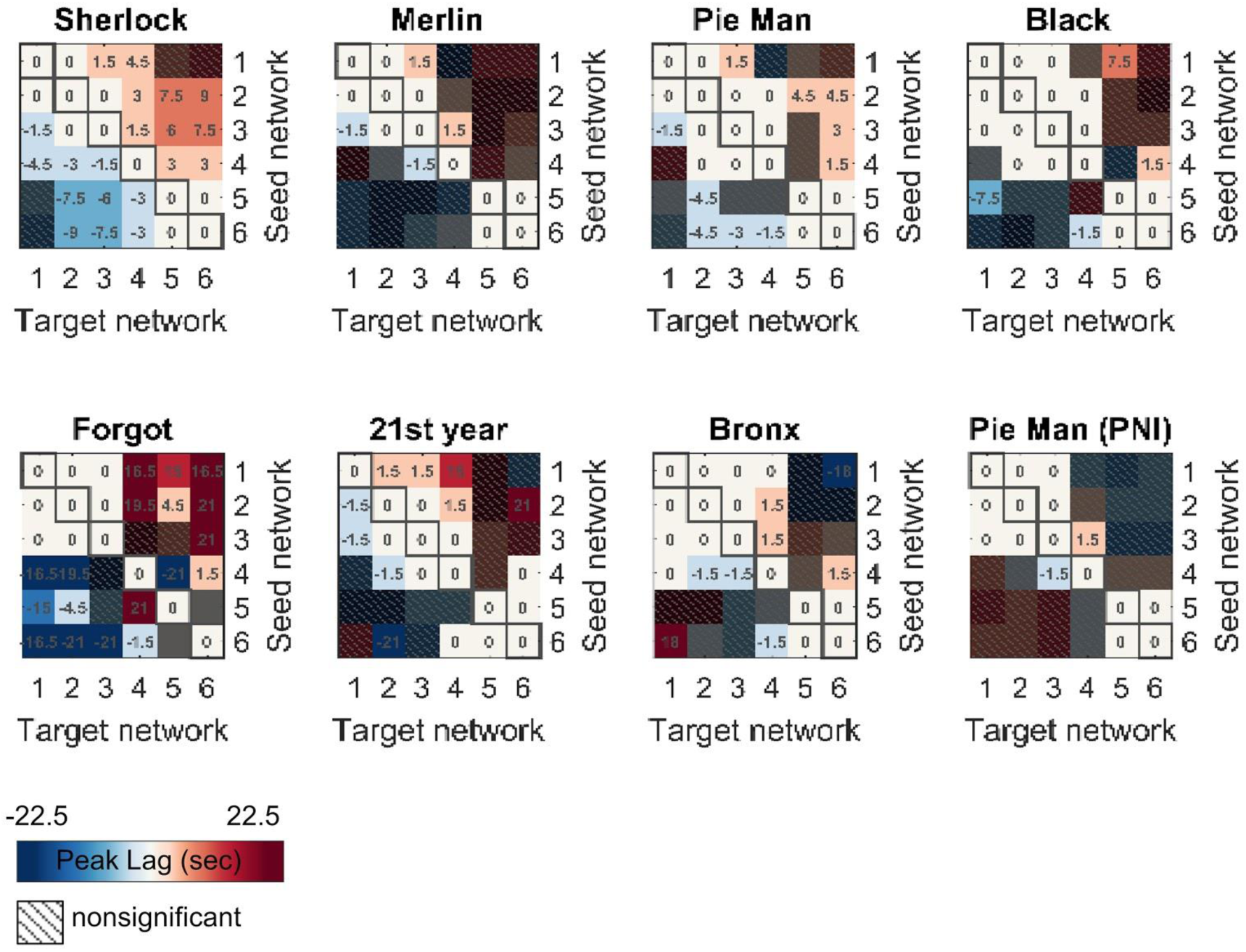
The network × network peak lag matrix based on the lag-ISFC in each individual story (p < .05, FDR corrected).

**Fig. S6.**
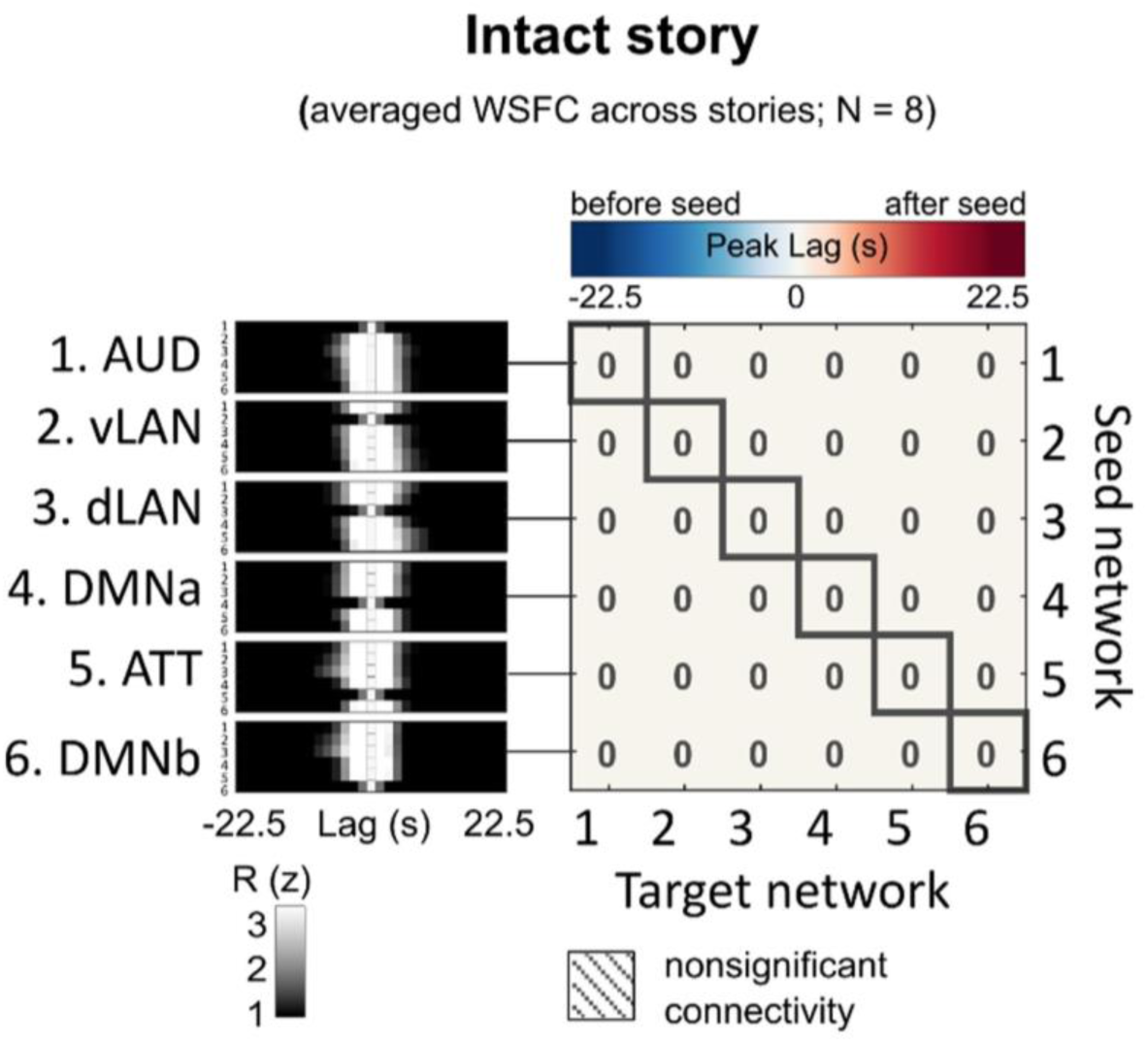
The network × network peak lag matrix based on the averaged lag-WSFC across eight stories (p < .05, FDR corrected).

**Fig. S7.**
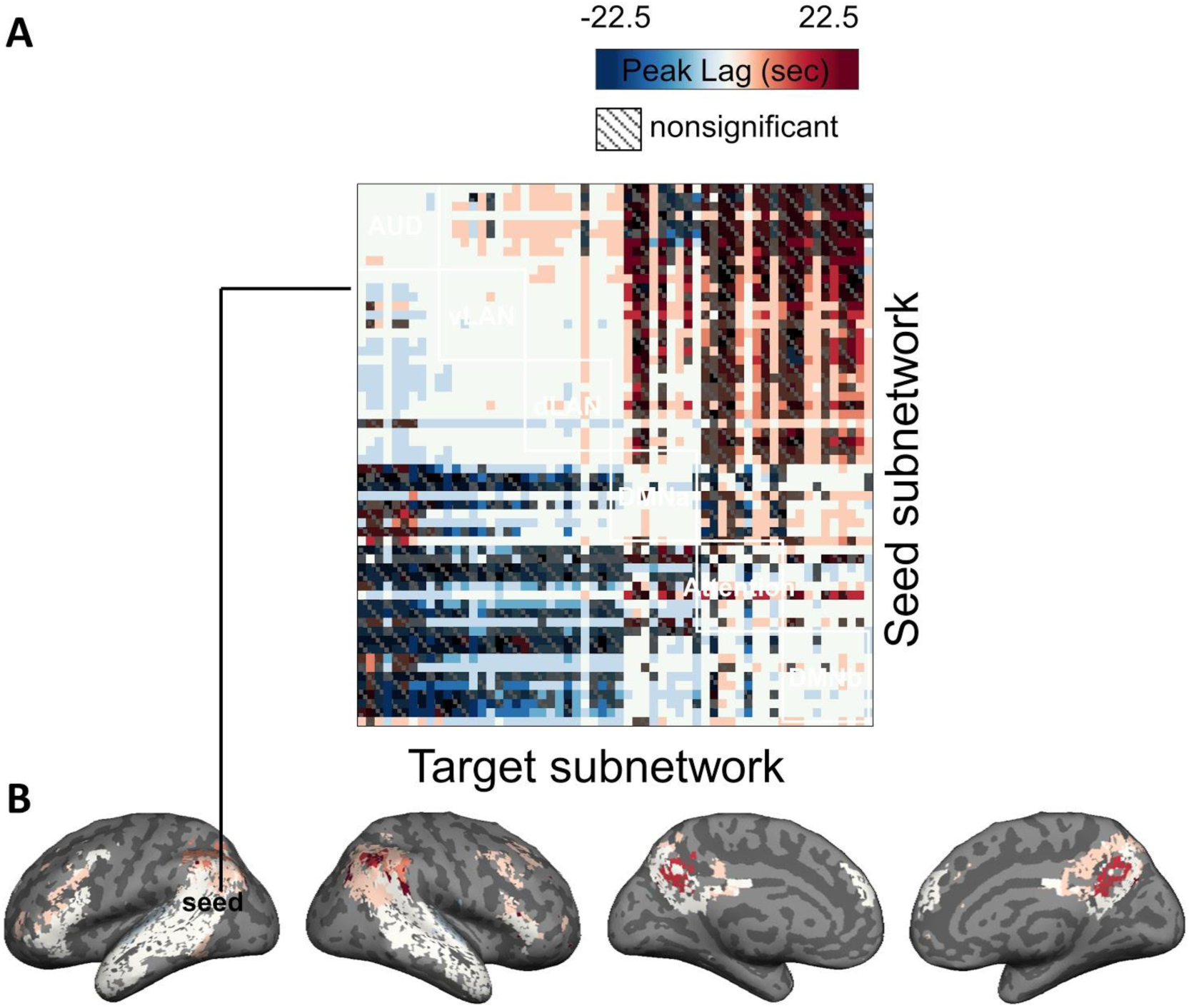
Subnetwork × subnetwork peak lags based on the averaged lag-ISFC across eight stories (p < .05, FDR corrected). The subnetworks were created by dividing each of the six main functional networks (SI Appendix, Fig. S1) into 10 subnetworks, applying k-means clustering to resting-state WSFC (k = 10 within each network). (A) The peak lag matrix. (B) The brain map of significant peak lags between one seed subnetwork (posterior superior/middle temporal gyrus) and all the sixty subnetworks.

**Fig. S8.**
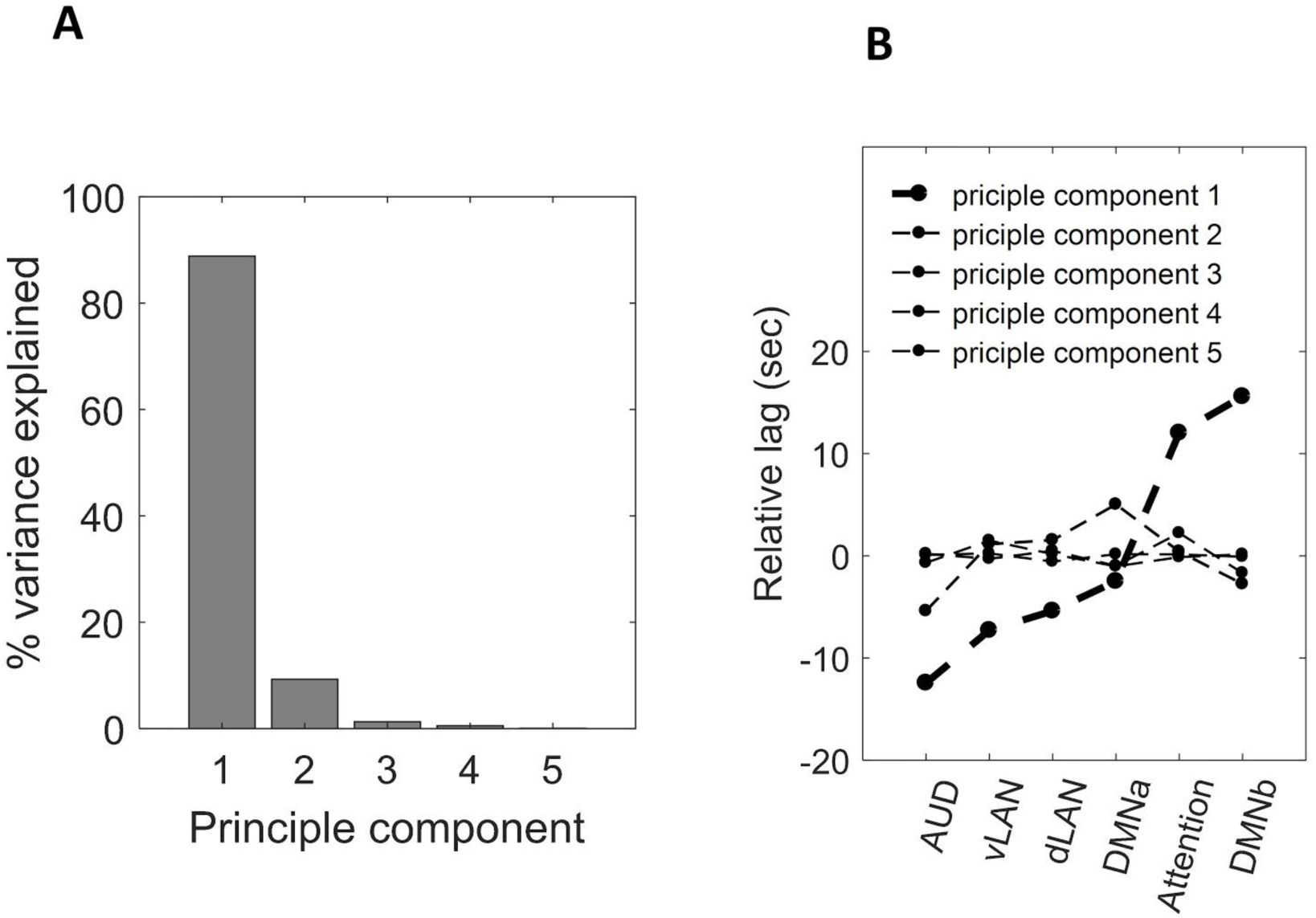
Principal component analysis of the inter-network lag matrix across eight stories (Fig. 3A). (A) The percentage of variance explained by each principal component. (B) Relative-lag values from each principal component. Line thickness indicates the percentage of variance explained by that component.

**Fig. S9.**
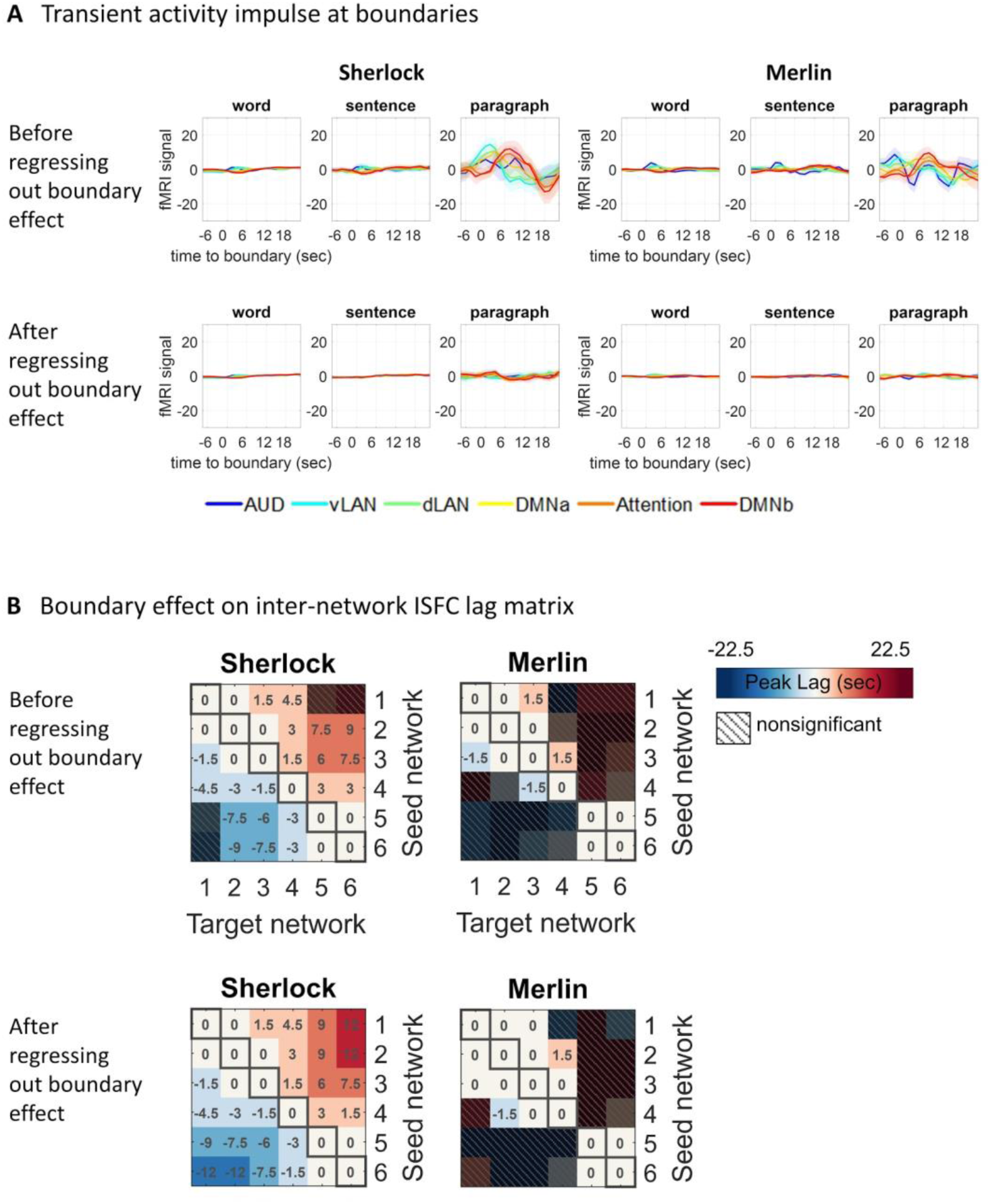
Boundary effect on the network x network peak lag matrix across stories. (A) The fMRI signals around word, sentence, and paragraph boundaries before and after regressing out the boundary effects. Shaded areas indicate 95% confidence intervals across subjects. (B) The peak lag matrix before and after regressing out the boundary effects (p < .05, FDR corrected).

**Fig. S10.**
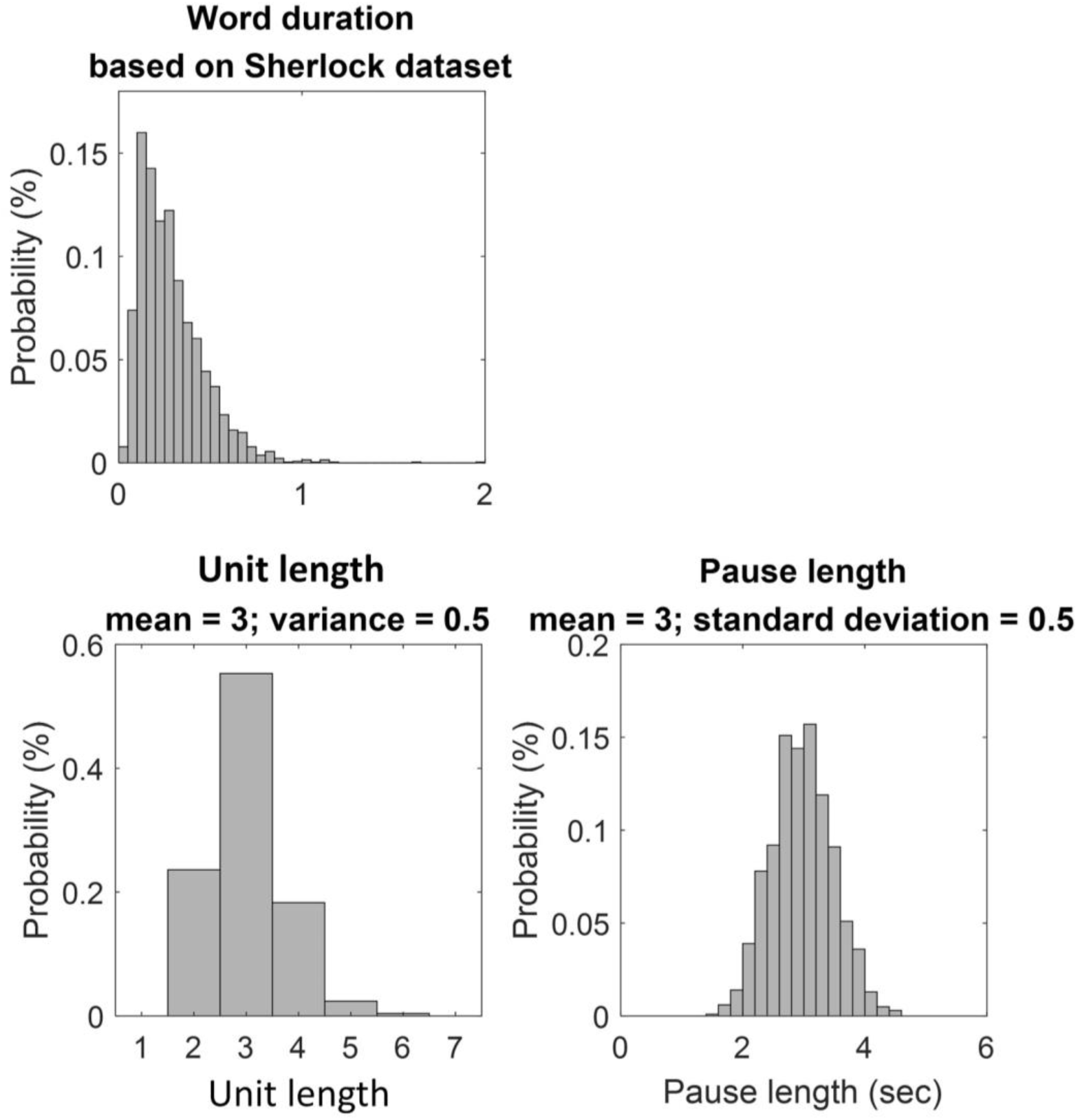
The distributions of word duration, unit length, and pause length with the simulation parameters described in Table S1.

**Fig. S11.**
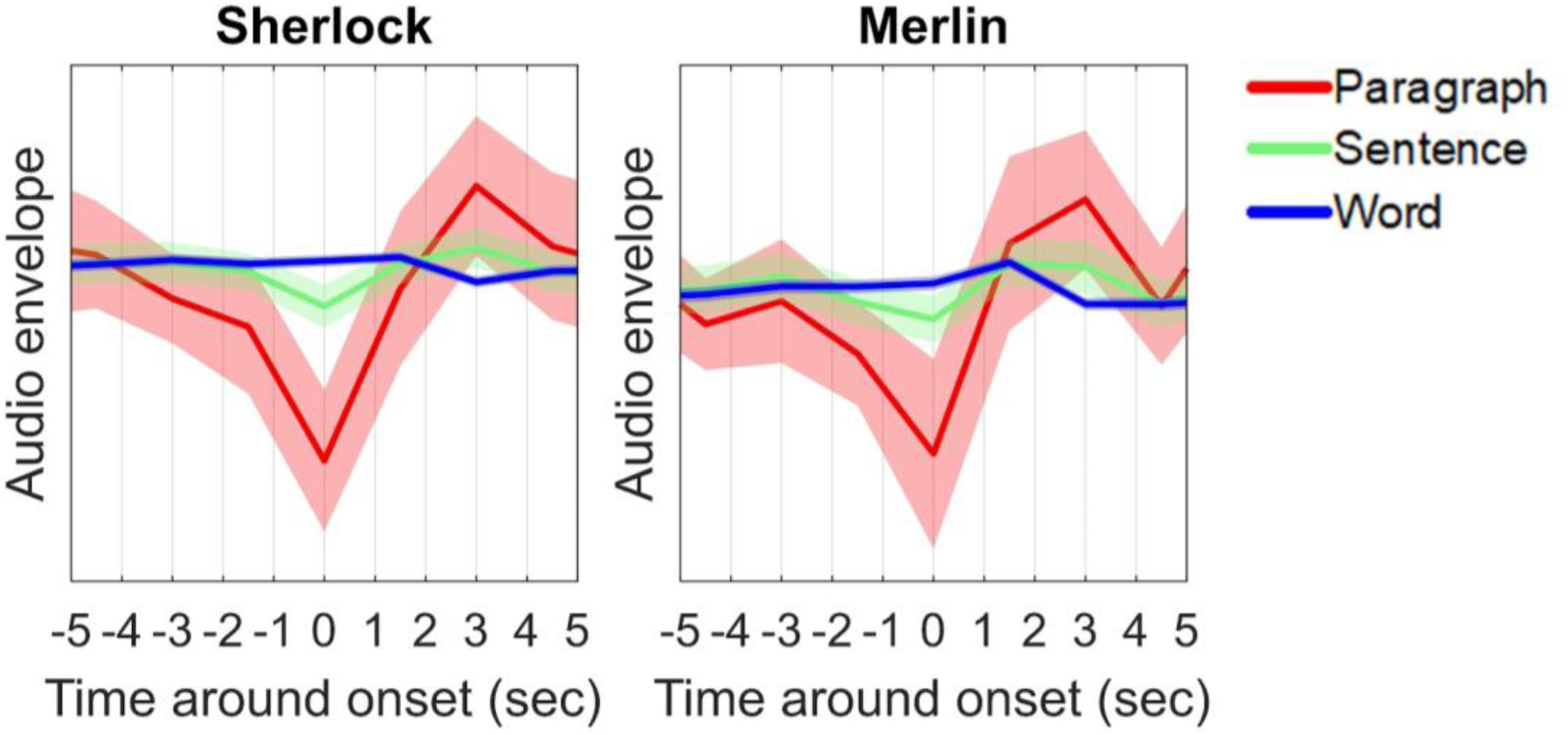
The silent pause between paragraphs shown in real spoken stories. Shaded areas indicate 95% confidence intervals.

**Fig. S12.**
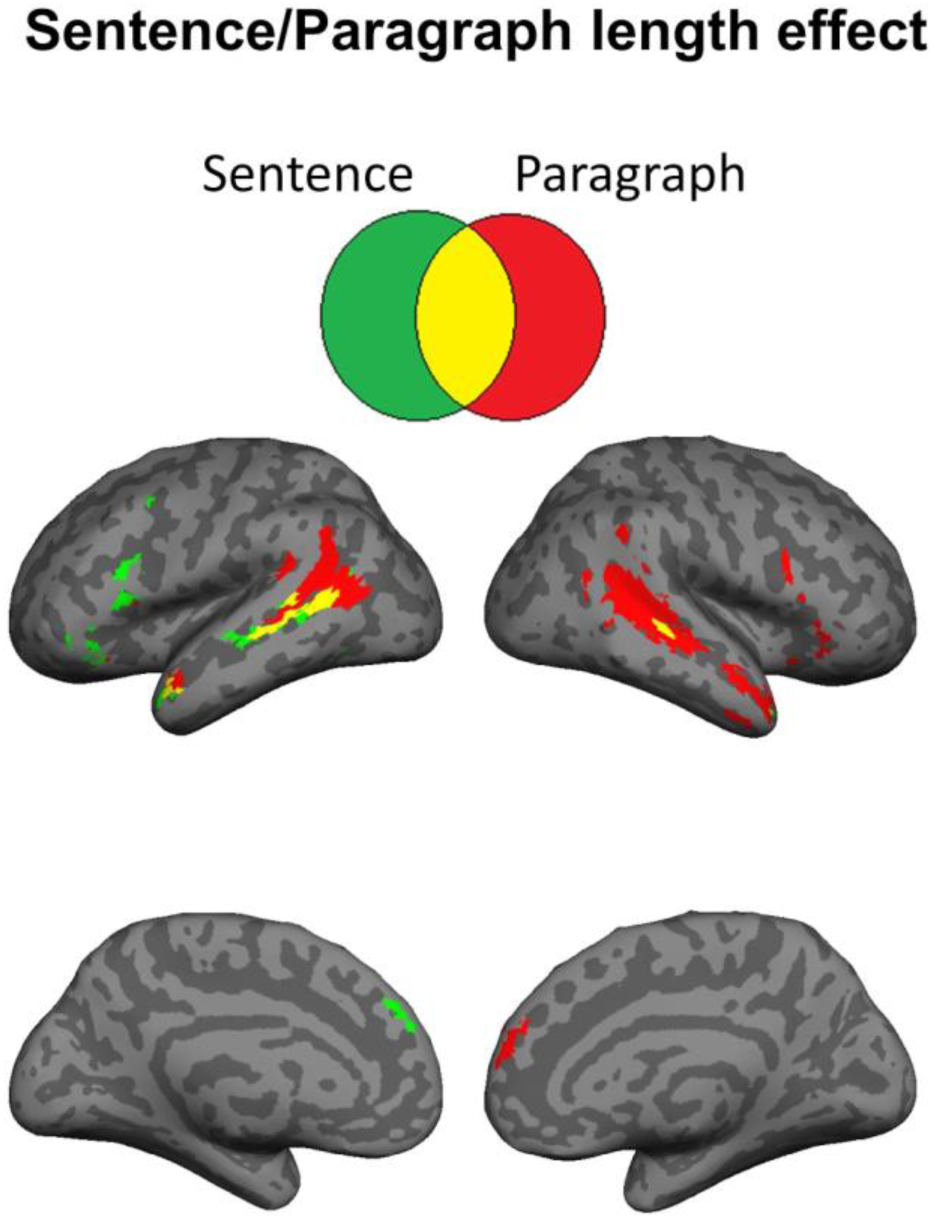
Sentence and paragraph length effects in two time-stamped stories (“Sherlock” & “Merlin”) (p < .005, uncorrected). Significant length effect indicates activation accumulation from the start toward the end of sentences or paragraphs.

**Fig. S13.**
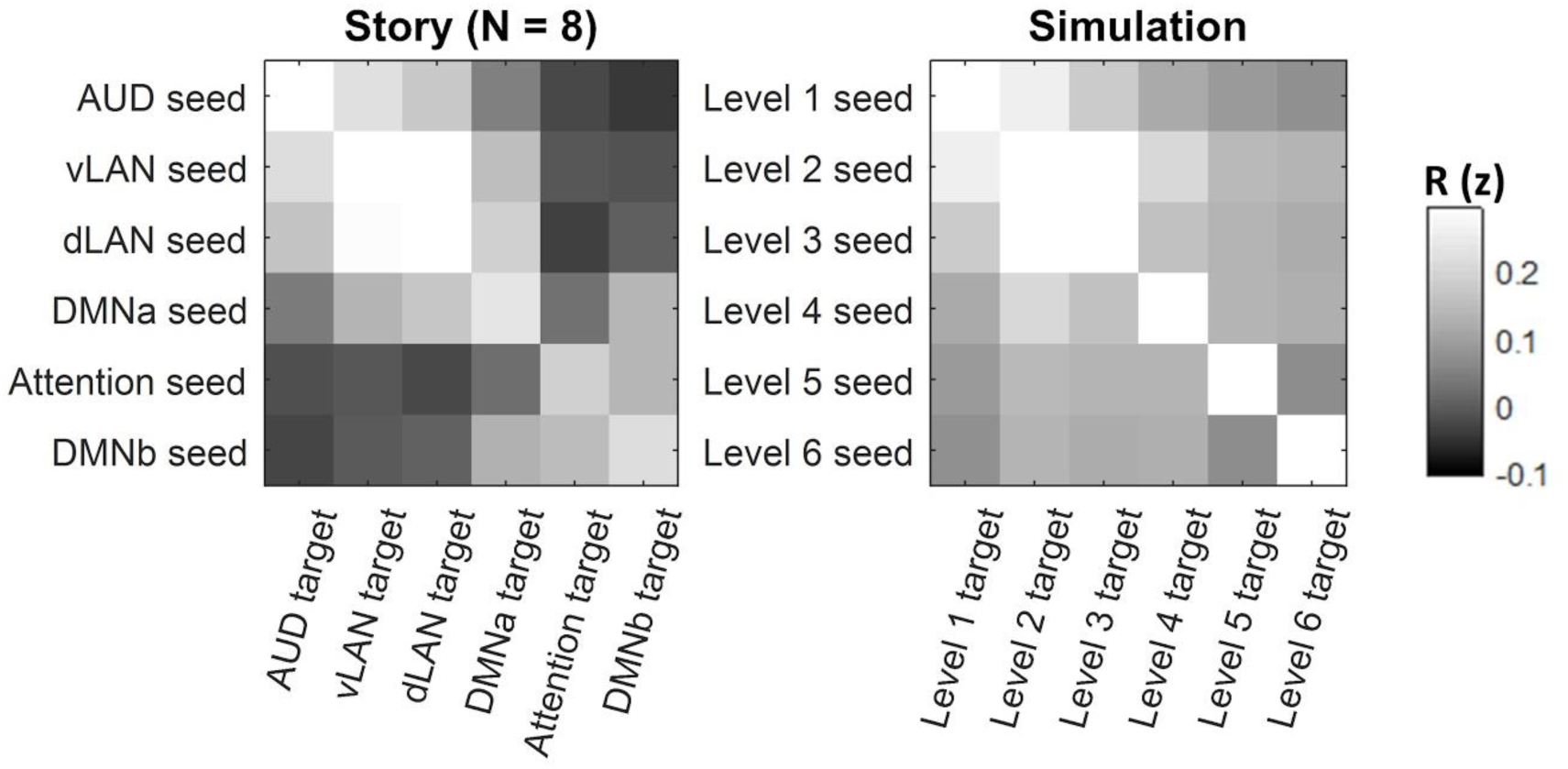
ISFC matrices at lag 0 in real and simulated stories (the same simulation parameters as in Table S1).

**Fig. S14.**
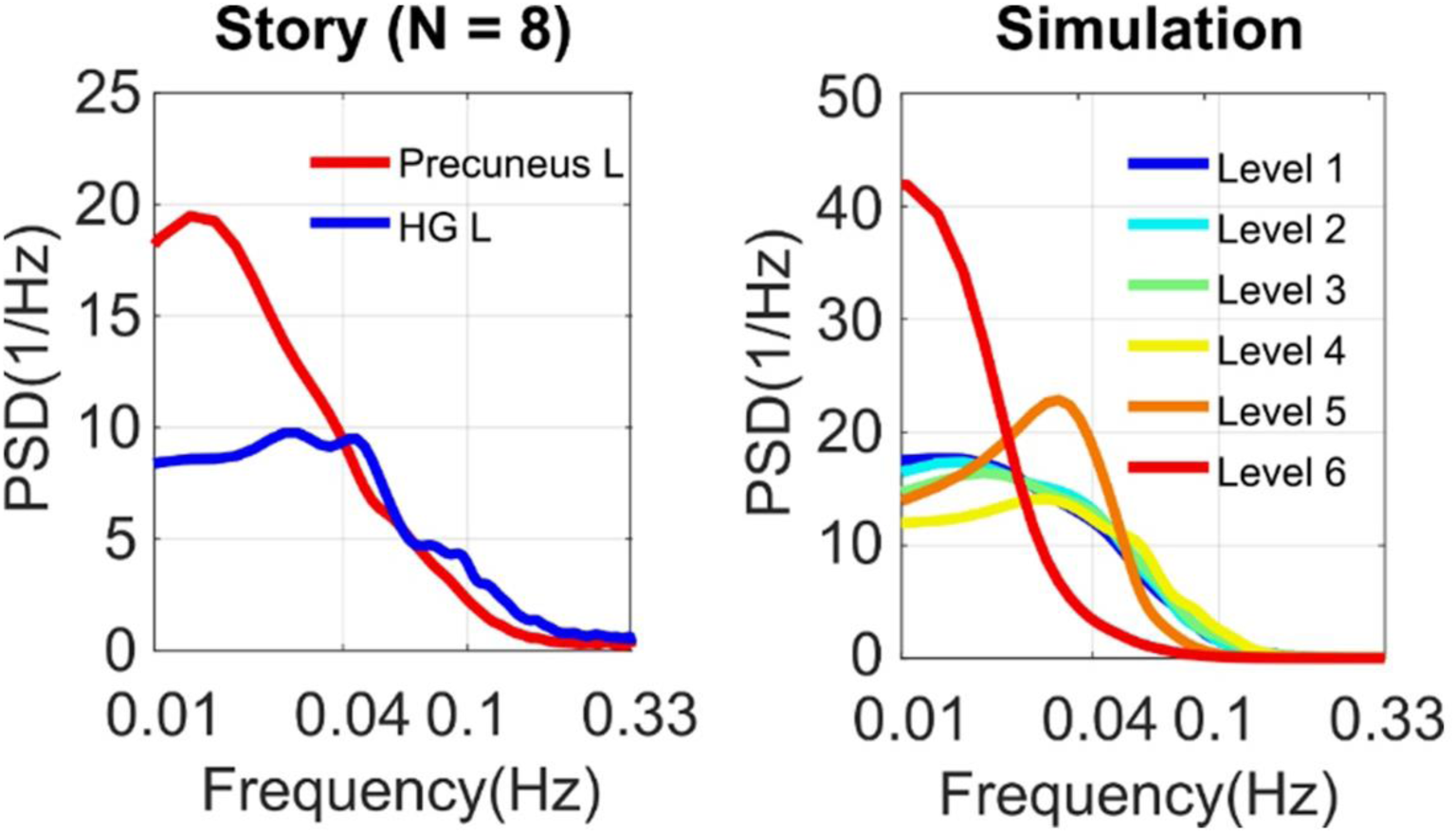
Power spectral densities of real (left) and simulated (right, the same parameter set as Table S1) BOLD responses to stories. PSD of the actual BOLD data exhibited stronger low-frequency fluctuations at regions with longer temporal receptive windows. Simulated BOLD responses show a similar pattern.

**Fig. S15.**
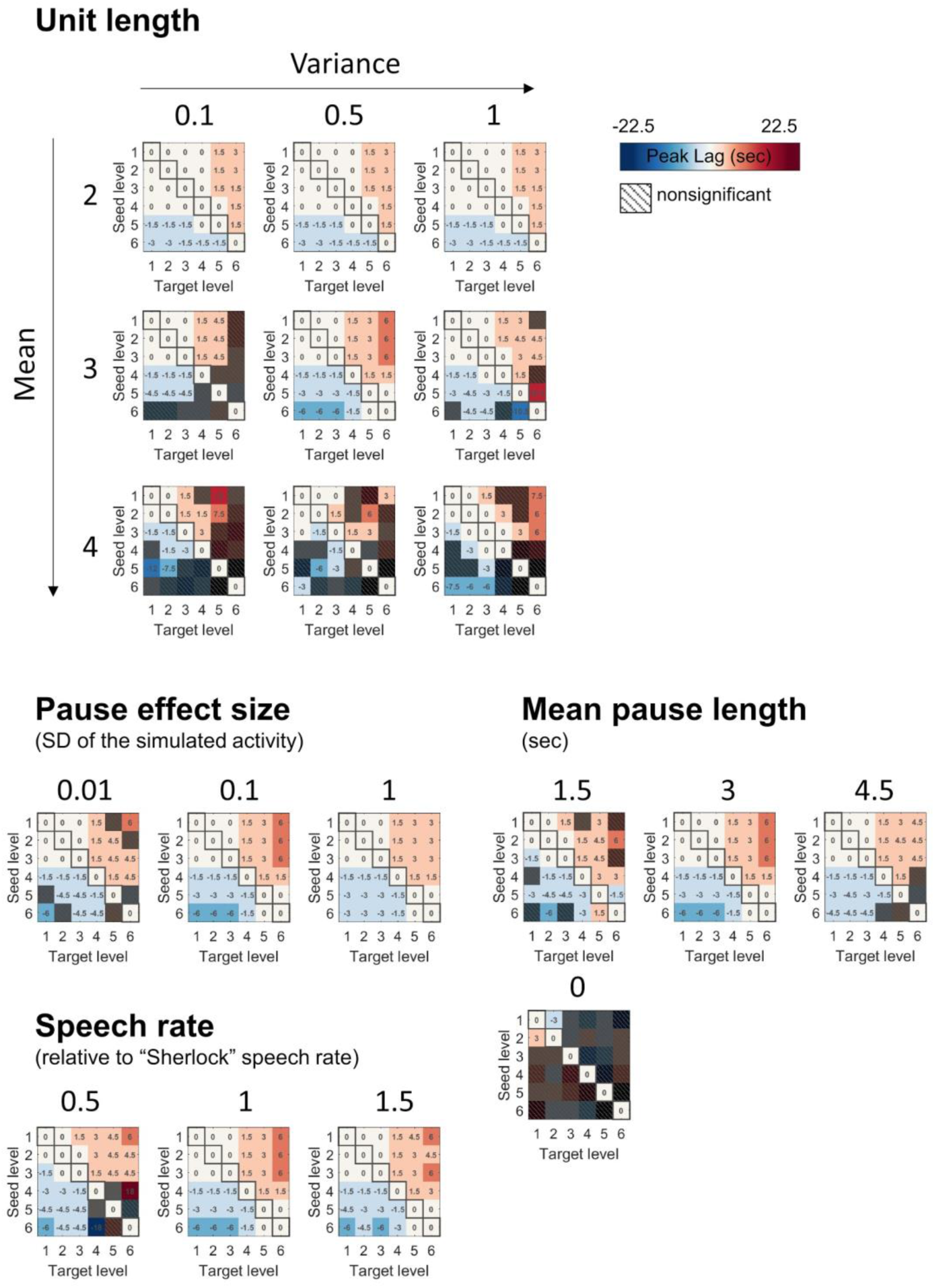
Robust lag gradient within the parameter space bound by natural speech (the same parameters as in Table S1 unless otherwise indicated) (p < .05, FDR correction).

**Table S1.** A set of exemplar stimulation parameters motivated by a spoken story (“Sherlock”). SD: standard deviation.

| Exemplar simulation parameters |  |
| --- | --- |
| speech rate (relative to “Sherlock”) | 1 |
| mean unit length | 3 |
| unit length variance | 0.5 |
| temporal integration function | linearly increasing |
| mean pause length | 3 sec |
| pause length effect size<br>(in SD of the simulated activity) | 0.1 |

## Notes

**Competing Interest Statement:** The authors declare no competing financial interests.

### Competing Interest Statement

The authors have declared no competing interest.

